# *HnRNPAB*-targeted antisense oligonucleotides ameliorate tau pathology and cognitive deficits by modulating Alzheimer’s disease-associated alternative splicing

**DOI:** 10.64898/2026.07.02.736008

**Authors:** Tao Jiang, Xingjun Xu, Yaqiong Yu, Caixin Zhu, Chao Ma, Yu Tang, Lingxia Min, Mingliang Tan, Jialin Bai, Zhou Feng, Jingming Hou

## Abstract

Aberrant alternative splicing alters multiple disease-associated splicing events, including MAPT exon 10 inclusion linked to tau pathology, and thereby impacts Alzheimer’s disease (AD) progression. Heterogeneous nuclear ribonucleoprotein AB (hnRNPAB) functions as a splicing regulator, yet its roles in AD remain poorly defined. Here, we identified AB332, the full-length isoform of hnRNPAB, as a novel repressor of *MAPT* exon 10 inclusion. AB332 binds a conserved AAUAU motif and recruits RBMX and RBMXL1 through its RRM and glycine-rich domains to assemble a complex. Beyond *MAPT*, AB332 modulates a network of splicing events in multiple AD-associated genes, including *STAG2, ApoER2, MCL-1, PICALM*, and *zDHHC7*. Importantly, the exon 7-skipping isoform AB285 lacks these activities. Transcriptomic analysis of post-mortem brain tissues from AD patients revealed reduced expression of *hnRNPAB*, while selective downregulation of AB332 was confirmed in the hippocampus of 3xTg AD mice. We performed a systematic antisense oligonucleotide-tiling screen targeting *hnRNPAB* exon 7 and its flanking intronic regions, identifying ASO30 that specifically upregulated AB332 expression. In 3xTg AD mice, ASO30 restored hippocampal AB332 levels, reduced pathological tau phosphorylation, and rescued cognitive deficits. These findings define a molecular mechanism by which AB332 regulates AD-associated splicing events, thereby providing a therapeutic strategy targeting *hnRNPAB* for AD.

**GRAPHICAL ABSTRACT:** 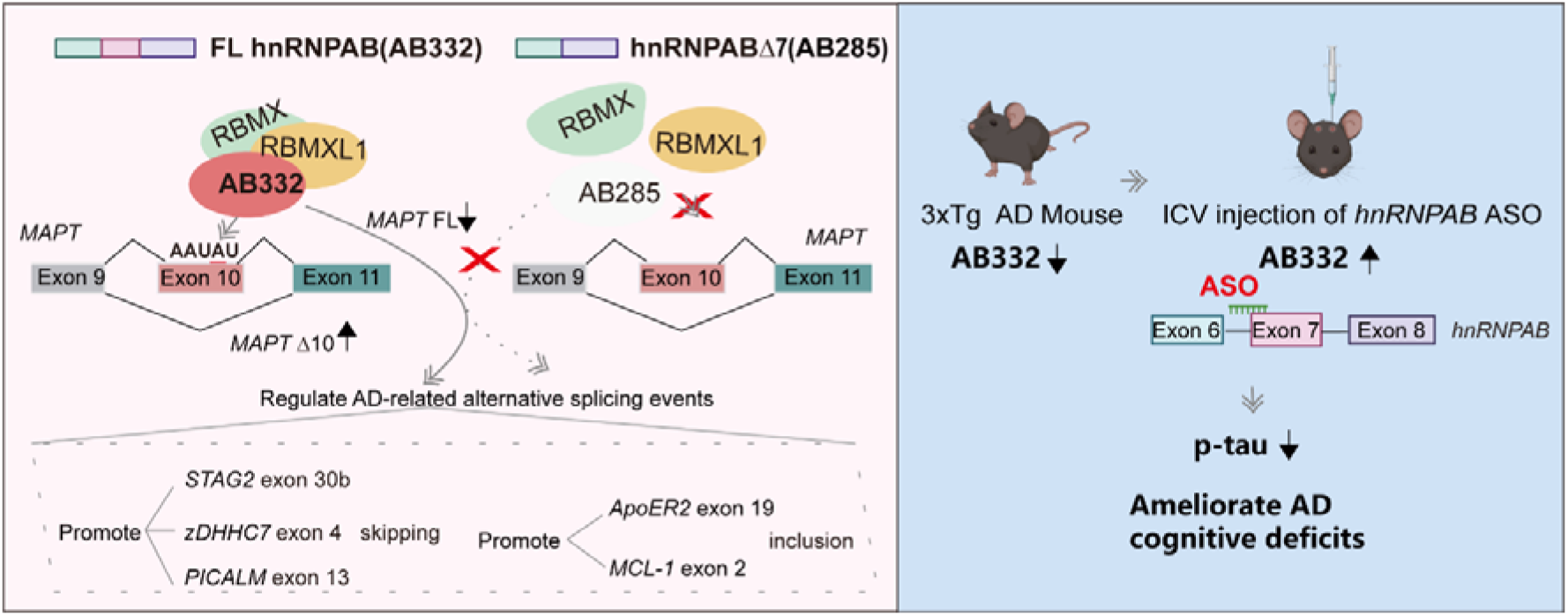

## INTRODUCTION

Alternative splicing serves as a pivotal post-transcriptional process that generates multiple distinct protein isoforms from a single pre-mRNA through mechanisms such as exon skipping and intron retention (1). Given its occurrence in over 90% of human genes, alternative splicing also serves as a major source of functional complexity in higher eukaryotes (2). This is particularly evident in the brain, which displays one of the highest prevalences of alternative splicing events among human tissues (3,4). Dysregulation of splicing regulatory elements or splicing factors can lead to aberrant alternative splicing patterns, which are critically involved in the neurodegenerative diseases (5,6).

Accumulating studies have established alternative splicing dysregulation as a vital factor affecting pathological progression in Alzheimer’s disease (AD), with its severity closely correlated with clinical disease stages (7). The typical pathological hallmarks of AD include amyloid plaques formed by β-amyloid (Aβ) deposition and neurofibrillary tangles induced by hyperphosphorylated tau protein (8). Among these, tau pathology shows a stronger association with dementia severity relative to Aβ deposition (9). The microtubule-associated protein tau (*MAPT*) gene, which encodes the tau protein, is one of the most representative core targets among genes with aberrant alternative splicing in AD. *MAPT* exon 10 has attracted extensive attention, as its alternative splicing generates two tau protein isoforms: 3-repeat tau (3R-tau, with three microtubule-binding repeats) and 4-repeat tau (4R-tau, with four such repeats) (10). In AD patients, the dysregulated 3R-tau:4R-tau ratio leads to increased tau phosphorylation (11). This isoform imbalance and its pathological consequences can be reversed by small-molecule splicing modulators that promote *MAPT* exon 10 skipping, thus reducing 4R-tau expression and hyperphosphorylated tau levels (12). Moreover, forebrain organoids infected with herpes simplex virus type 1 (HSV-1) exhibit enhanced inclusion of *MAPT* exon 10, elevating the 4R-tau ratio and recapitulating AD-like pathologies (13). These findings highlight the critical role of *MAPT* exon 10 splicing in AD. Beyond *MAPT*, transcriptomic studies consistently identify aberrant splicing in other AD-related genes—such as *APP, PICALM, CLU*, and *PTGDS*—supporting a network-based view of AD pathogenesis (7,14-17).

This multi-target landscape suggests that interventions focusing on single genes may be insufficient to reverse pathology(18), shifting therapeutic interest toward splicing regulators. Targeting these factors, which coordinate entire splicing networks, offers a promising “one-target, multi-effect” strategy. Consistently, proteomic analyses of AD brains reveal widespread alterations in splicing-related RNA-binding proteins (19-21). For instance, overexpression of the splicing factor SFA-1 alleviates age-related global splicing defects and extends lifespan (22). Similarly, diminished NOVA1/2 protein levels in AD correlate with disease-relevant, aging-dependent splicing changes (23). Heterogeneous nuclear ribonucleoprotein (hnRNP) family members further exemplify this therapeutic axis: downregulation of hnRNPA1 (a regulator of *APP, RAGE*, and *MAPT* splicing) exacerbates pathogenesis (24-27); the neuroprotective hnRNPL-DL isoform modulates a network of synaptic genes including *CAMKV* (28). Therefore, these findings underscore the broad potential of correcting splicing regulator dysfunction in AD.

HnRNPAB is a canonical splicing regulator within the hnRNP family. While its roles in cancers are well-established (29-32), preliminary evidence implicates hnRNPAB in neurobiological processes. This includes its binding to mRNAs crucial for axonal guidance and synaptogenesis (33), as well as its promotion of recovery after neural injury through pathways such as NF-*κ*B/Lcn2 (34). Nevertheless, its function in AD, remains poorly understood. In this study, we report that the full length hnRNPAB (AB332) forms a complex with RBMX/XL1 to specifically inhibit *MAPT* exon 10 splicing through recognition of a conserved AAUAU motif. Beyond *MAPT*, AB332 also modulates the splicing of multiple AD-associated genes, whereas the exon 7-skipped isoform lacks this splicing regulatory activity.

Hippocampal expression of AB332 is significantly reduced in 3xTg AD mice. Restoring its levels using antisense oligonucleotides (ASOs) that promote *hnRNPAB* exon 7 inclusion effectively rescues cognitive deficits in this AD model. Together, these findings establish hnRNPAB-directed splicing correction as a novel therapeutic strategy for AD, highlighting the potential of targeting splicing networks to modify disease pathogenesis.

## MATERIAL AND METHODS

### Plasmid construction

The *MAPT* minigene construct used in this study was pCI-MAPT, as previously described (35). Briefly, this minigene contains 265-nt exon 9, a truncated 388-nt intron 9, 93-nt exon 10, 429-nt intron 10, and the first 82 nt of exon 11. Minigene mutants were generated via site-directed mutagenesis using partially overlapping primer pairs. To construct the pCI-MAPT-boxB plasmid, the AATAT sequence (positions +64 to +68) in exon 10 of the *MAPT* minigene was replaced with a 19-nt boxB hairpin sequence (5’-GGGCCCTGAAGAAGGGCCC-3’). All protein expression plasmids were constructed in either the pCGT7 or FLAG vector, with expressed proteins carrying an N-terminal T7 or FLAG tag, respectively. Detailed sequences of all primers are provided in Table S1.

### Cell culture and transfection

BV2, HeLa, SK-N-SH and N2A cell lines were purchased from Procell Life Science & Technology Co., Ltd. (Wuhan, China). Among these, BV2 and HeLa cells were cultured in Dulbecco’s modified Eagle’s medium (DMEM; GIBCO, Grand Island, NY, USA), while SK-N-SH and N2A cells were maintained in Minimum Essential Medium (MEM; GIBCO, Grand Island, NY, USA) supplemented with non-essential amino acids (NEAA; GIBCO). All cell lines were grown in medium containing 10% (v/v) fetal bovine serum (FBS) and antibiotics (100 U/ml penicillin and 100 μg/ml streptomycin) at 37°C in a humidified 5% CO_2_ atmosphere. To analyze the effect of protein overexpression on wild-type (WT) and mutant *MAPT* minigenes, 1.5×10^5^ cells per well were seeded into 12-well plates containing DMEM supplemented with 10% FBS. The following day, 500 ng of each minigene plasmid was co-transfected with 300 ng of each protein-expression plasmid or Vctr into the cells using ExFect Transfection Reagent (Vazyme, Nanjing, China). For hnRNPAB knockdown, si-hnRNPAB (sense: 5’-UGGAAGCAAGUGUGAGAUCAATT-3’) was used, with siNC (sense: 5’-UUCUCCGAACGUGUCACGU-3’) serving as the negative control. Each siRNA was transfected alone or co-transfected with a minigene plasmid, using the same ExFect Transfection Reagent for consistency. After incubation at 37°C for approximately 6 hours, the transfection medium was removed and replaced with complete medium. All ASOs used in this study were uniformly modified with 2’-O-methoxyethyl groups, a phosphorothioate backbone, and 5-methylcytosines throughout the sequence. These ASOs were synthesized and purchased from Hippo Biotechnology Co., Ltd. Detailed sequences of all ASOs are provided in Table S2. For ASO screening assays, individual ASOs were transfected into HeLa cells using the CALNP™ RNAi in vitro transfection reagent (Duona Pharmaceutical Co., Ltd.) following the manufacturer’s recommended protocol.

### Semi-quantitative RT-PCR and qRT-PCR

Cultured cells were collected ∼36 hrs post-transfection and total RNA was isolated with Trizol reagent (Vazyme, Nanjing, Jiangsu, China), followed by first-strand cDNA synthesis with 500 ng of each RNA sample using oligo (dT)18 and M-MLV reverse transcriptase (Vazyme) in a 10 μL reaction. For minigenes, splicing products were amplified using 28 PCR cycles with forward primer MAPT-F (5’-CCACTCCCAGTTCAATTACAGC-3’) and MAPT-R (5’-Cy5-TAATGAGCCACACTTGGAGGTC-3’) (35). while for the endogenous *MAPT*, we used 35 cycles with forward primer EMAPT-F (5’-CCATGCCAGACCTGAAGAAT-3’) and reverse primer EMAPT-R (5’-TGCTCAGGTCAACTGGTTTG-3’). Primer sequences for validating endogenous alternative splicing of *hnRNPAB, ApoER2, TMPO, zDHHC7, MCL-1, STAG2*, and *PICALM* are listed in Table S3. Cy5-labeled PCR products were separated on 6% native polyacrylamide gels, followed by fluorescence imaging with Biorad ChemicDoc MP Imaging system and signals were quantitated by Biorad Image Lab (Biorad, Hercules, CA). Exon inclusion was expressed as a percentage of the total amount of spliced mRNA. qRT-PCR were performed using a SYBR qRT-PCR kit (Vazyme, Nanjing, Jiangsu, China). GAPDH was used as reference gene. cDNAs were amplified with 40 cycles using specific primers and fold changes were calculated using the 2-ΔΔCt method. The sequences of all primers used for RT-qPCR are listed in Table S3.

### Western blot

Cultured cells and homogenized mouse tissues were lysed in RIPA buffer (Beyotime, Shanghai, China) supplemented with protease inhibitor cocktail and/or phosphatase inhibitor cocktail (ABclonal, Wuhan, China). Protein samples were separated by 10–12% SDS-PAGE and then transferred onto PVDF membranes (Immobilon-P, Millipore, MA, USA). The membranes were incubated with primary antibodies at 4 ºC overnight, followed by incubation with IRDye 680RD goat anti-mouse or anti-rabbit secondary antibodies (LI-COR Biosciences, Lincoln, NE, USA). Detailed information of antibodies is provided in Table S4. Signals were detected using a Bio-Rad imaging system with the IRDye 680RD channel for visualization. Protein band intensities were quantified, normalized to the corresponding loading control, and further normalized to the control group.

### RNA pull-down

RNA pull-down was performed as previously reported(35). biotin-labeled WT RNA (5’-UCCUUAAAUUAA-3’) and mutant RNA (5’-UCCGGAAAUUAA-3’) were from General Biol. 500 pmol RNA was incubated with 60 μL pre-washed streptavidin agarose beads (Sigma-Aldrich) at 4°C for 2 h with rotation, respectively. The RNA-bead complexes were then incubated with HeLa cell nuclear extract in binding buffer (20 mM HEPES pH7.8, 100 mM KCl, 0.1-0.2 mM EDTA, 1 mM DTT, 1 mM PMSF, 0.05% Triton X, 0.25 μg/μL yeast tRNA, and protease inhibitor cocktail) at 4°C for 2 h. Beads were washed three times with buffer (150 mM KCl) and bound proteins were eluted with Laemmli buffer, heated at 95°C for 10 min for subsequent analysis.

### IP/Co-IP

FLAG-AB332 and FLAG-AB285 expression vectors were separately overexpressed in HeLa cells. Total proteins were extracted, and the cell lysate was incubated with magnetic agarose beads covalently coupled to anti-FLAG nanobodies for 3 h. The bound proteins were eluted and the anti-FLAG beads were resuspended. Samples were separated by SDS-PAGE, stained with Coomassie brilliant blue, and differential protein bands were selected for mass spectrometry analysis.

### Isoform sequencing and alternative splicing analysis

For Iso-Seq, total RNA was extracted from HeLa cells overexpressing AB332 and Vctr using the Direct-zol RNA Miniprep Kit (R2050, ZYMO Research, USA) according to the manufacturer’s instructions. Referring to the Isoform Sequencing protocol (Part Number PN 100-092-800-03) released by Pacific Biosciences, the sequencing library was constructed with the Clontech SMARTer PCR cDNA Synthesis Kit (Catalog# 634938, Clontech Laboratories, CA, USA). Libraries were then sequenced on the PacBio Sequel third-generation sequencing platform (Pacific Biosciences, USA), which is specifically designed for Iso-Seq and enables full-length transcriptome sequencing without the need for fragmentation. Sequencing reads were aligned to the human genome (Homo sapiens. GRCh38.p14 assembly). Alternative splicing analysis was performed using SUPPA software to calculate the percent spliced in (PSI) values. The change in PSI (ΔPSI) was defined as the difference between the PSI value of the AB332 overexpression group (PSI_AB332 OE_) and that of the control group (PSI_Vctr_): a positive ΔPSI (ΔPSI > 0) indicates an increased inclusion level of the target alternative splicing event, while a negative ΔPSI (ΔPSI < 0) denotes a decreased inclusion level.

### Enhanced crosslinking and immunoprecipitation sequencing

AB332 plasmid was transfected into HeLa cells in T75 flasks. At 48 h post-transfection, cells were UV-crosslinked on ice with a Stratalinker crosslinker (Stratagene) using 254 nm UV (600 mJ/cm^2^) to convert protein-RNA ionic bonds to covalent bonds. Cells were lysed with CLIP lysis buffer, RNA was enriched for immunoprecipitation as previously reported, and separate eCLIP and input libraries were constructed for each sample. 400 μl nuclear extract was mixed with 5 μg AB332-specific antibody and incubated overnight at 4°C for immunoprecipitation to enrich AB332-RNA complexes. The next day, Protein A/G beads were added for 4 h incubation, followed by two washes with IP buffer to remove non-specific bindings. Purified products were separated by 4%–12% Bis-tris gel electrophoresis, transferred to a nitrocellulose membrane, and regions containing AB332 were excised to extract RNA-RNP complexes for dephosphorylation. A single-stranded RNA adapter was ligated to RNA 3’ end, and RNA 5’ end was ^32^P-labeled with T4 polynucleotide kinase (3’ phosphatase minus). AB332-RNA complexes were recovered, proteins were removed by proteinase K and urea treatment, and target RNA was purified for reverse transcription to generate first-strand cDNA. After removing residual RNA and dNTPs, a DNA adapter was ligated to cDNA 3’ end, followed by PCR amplification and gel-based size selection to construct the AB332 eCLIP library.

### Animal Experiments and ASO Administration

C57BL/6J and 3xTg-AD mice were obtained from Jinzhihe Biotechnology (Guangzhou, China). All animal experimental protocols were approved by the Ethics Committee of Third Military Medical University (Chongqing, China). All experimental mice were housed in a standardized environment with a 12-h light/dark cycle, maintained at 23°C and 50-60% relative humidity, with free access to standard rodent chow and water. To initially detect AB332 expression in AD, four 12-month-old female 3×Tg AD mice and four age-matched female WT mice were used. For testing the effect of ASO30 on exon 7 splicing of *hnRNPAB*, 6-week-old female C57BL/6J mice were employed. To evaluate the impacts of ASO30 on cognitive function and AD pathology, 9-month-old female 3×Tg AD mice were selected. ASO30 was administered via a single left lateral ventricle injection at a dose of 100 ng per mouse. Mice were anesthetized and then fixed on a stereotaxic instrument, with ASO30 injected using a 10 *μ*L Hamilton syringe at predetermined coordinates. Specifically, 6-week-old mice received injections in the hippocampus at coordinates (relative to the lambda point): AP -0.3 mm, ML 1 mm, DV -2.3 mm; 7-month-old mice were injected at distinct hippocampal coordinates (relative to the lambda point): AP -0.5 mm, ML 1.3 mm, DV -2.6 mm. All mice were given analgesics during surgery and monitored throughout the surgical procedure as well as for 3 days postoperatively.

### Behavioural assays

#### Open field

Exploratory behavior was evaluated using a 40×40×40 cm opaque arena. Mice were acclimatized to the testing environment for 30 min, placed individually in the arena center, allowed 2 min adaptation, then tested for 10 min. Total ambulatory distance (cm) and center zone occupancy time (%) were automatically tracked and quantified by EthoVision XT (v11.5, Noldus).

#### Novel object recognition (NOR)

Recognition memory was assessed in a 40×40 cm open field arena. Twenty-four hours after environmental habituation, mice were placed in the arena center with two identical objects (5 cm away from the walls) and allowed to explore for 10 min (acquisition phase). After a 24-hour interval, one familiar object was replaced with a novel one, and mice were allowed to explore for 10 min (test phase). All exploration behaviors were video-recorded and automatically tracked using EthoVision XT software (v11.5, Noldus).

#### The Morris water maze

Spatial learning and memory were evaluated using a circular pool (120 cm diameter) filled with opaque water (23°C, 30 cm depth). Visual cues (geometric shapes) were placed around the pool. Prior to training, mice were allowed to swim freely in the pool for 2 min to habituate to the environment. The experiment lasted 5 days: over the first 4 days (training phase), mice underwent four daily trials (one from each quadrant, 60 min inter-trial interval) to locate a submerged annular platform (10 cm wide, 1 cm below water surface) within 60 s. Mice failing to find the platform within 60 s were guided to it and allowed to stay for 15 s. Latency to the platform was recorded using EthoVision XT software (v11.5, Noldus), with the daily mean latency calculated. On day 5 (probe trial), the platform was removed, and mice explored the pool freely for 60 s. Parameters including quadrant dwell time (%), number of platform crossings were collected and analyzed.

#### Statistical analysis

Data were analyzed using two-tailed Student’s t-test and one-way analysis of variance (ANOVA) followed by post hoc tests, with SPSS software (v27.0). All data are presented as mean ± standard deviation (SD). A *P* value < 0.05 was considered statistically significant.

## RESULTS

### HnRNPAB represses MAPT exon 10 splicing via the element downstream of exon 10

To elucidate the splicing regulatory function and underlying molecular mechanism of hnRNPAB, we utilized our previously validated *MAPT* minigene reporter system and co-expressed T7-tagged hnRNPAB with this minigene in HeLa cells (35). Fluorescent semi-quantitative RT-PCR analysis (36) revealed that hnRNPAB overexpression dramatically reduced *MAPT* exon 10 inclusion from 35% to 3% compared with the empty vector control (Vctr) (Fig. 1A-B). Western blotting with an anti-T7 antibody confirmed T7-hnRNPAB expression (Fig. 1C). SiRNA-mediated knockdown of hnRNPAB significantly increased *MAPT* exon 10 inclusion (Fig. 1D-F). Together, these data indicate hnRNPAB as a potent repressor of *MAPT* exon 10 splicing.

**Fig. 1.**
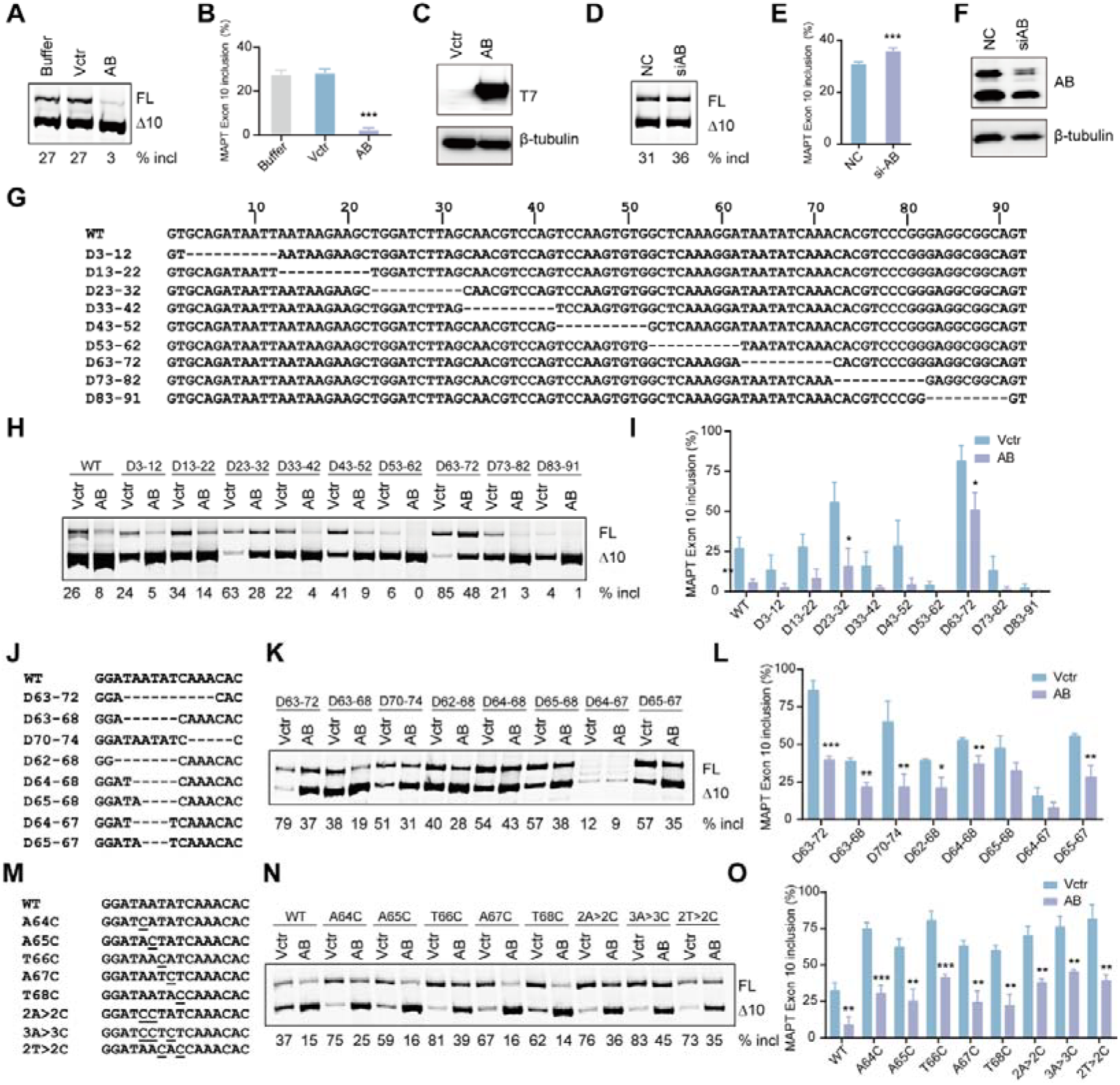
hnRNPAB-dependent *MAPT* exon 10 splicing inhibition via exonic element. **(A-C)** Effects of T7-tagged hnRNPAB overexpression on *MAPT* minigene splicing and quantification in HeLa cells. T7-tagged hnRNPAB expression was verified by Western blot (n = 3). **(D-F)** Effect of siRNA-mediated hnRNPAB knockdown on *MAPT* minigene splicing and quantification in HeLa cells (n = 4). Protein levels were examined by Western blot after siRNA treatment. **(G-I)** WT *MAPT* exon 10 sequence and the regions deleted (dashed lines) in the initial exonic deletion mutants. Splicing and quantitative analysis of WT and mutant *MAPT* minigenes upon T7-tagged hnRNPAB overexpression (n = 3). **(J-L)** Exon 10 mutants with further deletions spanning positions 62–73, and corresponding splicing patterns and quantification upon T7-tagged hnRNPAB overexpression (n = 3). **(M-O)** Substitution mutants were generated by targeting positions 64–68 of exon 10 (mutated bases underlined), with corresponding splicing patterns and quantification upon overexpression of T7-tagged hnRNPAB (n = 3). \**P* < 0.05, \*\**P* < 0.01, \*\*\**P* < 0.001 vs Vctr. \*\*\**P* < 0.001 vs NC.

To pinpoint the hnRNPAB-binding element, we generated a series of deletion mutants spanning exon 10 and flanking intronic regions (−50/+50) in the *MAPT* minigene (Fig. 1G and Fig. S1A), including 10 intronic and 9 exonic deletion mutants with intact exon-terminal dinucleotides to avoid disrupting splice sites. Following co-transfection with T7-hnRNPAB or Vctr, four intronic and two exonic deletions nearly abolished exon 10 inclusion (<10%), precluding further analysis; hnRNPAB overexpression strongly repressed exon 10 inclusion in most remaining mutants but showed markedly reduced activity in mutant D63–72 (Fig. 1H–I, S1B–C), indicating that the +63–72 region of exon 10 harbors a critical cis-element for hnRNPAB-mediated splicing silencing. We further refined this region using targeted deletions and found that removal of the 5-nt AAUAU motif (ΔD64–68) strongly impaired hnRNPAB-dependent repression (Fig. 1J–L). Systematic nucleotide substitution within this motif revealed that a single point mutation (U66C) was sufficient to disrupt hnRNPAB-mediated splicing inhibition (Fig. 1M–O). Collectively, our systematic mutagenesis studies identify the +64–68 AAUAU motif as a functional exonic splicing silencer (ESS) required for hnRNPAB-mediated repression of *MAPT* exon 10 inclusion.

### Specific binding of hnRNPAB to the *MAPT* exon 10 AAUAU element mediates splicing repression

To validate the mechanistic link between hnRNPAB-mediated splicing repression and AAUAU motif binding, we conducted RNA pull-down assays using biotinylated 12-nt RNA probes spanning exon 10 positions +59 to +72, with a T66C mutant as a non-targeting control. LC-MS analysis identified four RNA-binding proteins that were enriched in the WT sample, with peptide-spectrum matches (PSMs) ≥ 15 and a WT/mutant PSM ratio > 2: hnRNPD, hnRNPDL, hnRNPAB, and TIAL1 (Fig. 2A). To further confirm the specificity of hnRNPAB, we examined the effects of three additional RNA-binding proteins on *MAPT* minigene splicing. We found that hnRNPDL and TIAL1 significantly repressed *MAPT* exon 10 inclusion, whereas hnRNPD exhibited no regulatory effect (Fig. 2B-C). Western blot analysis showed significant enrichment of hnRNPAB in WT RNA–protein complexes relative to mutant controls, while hnRNPDL and TIAL1 exhibited no such preferential binding (Fig. 2D).

**Fig. 2.**
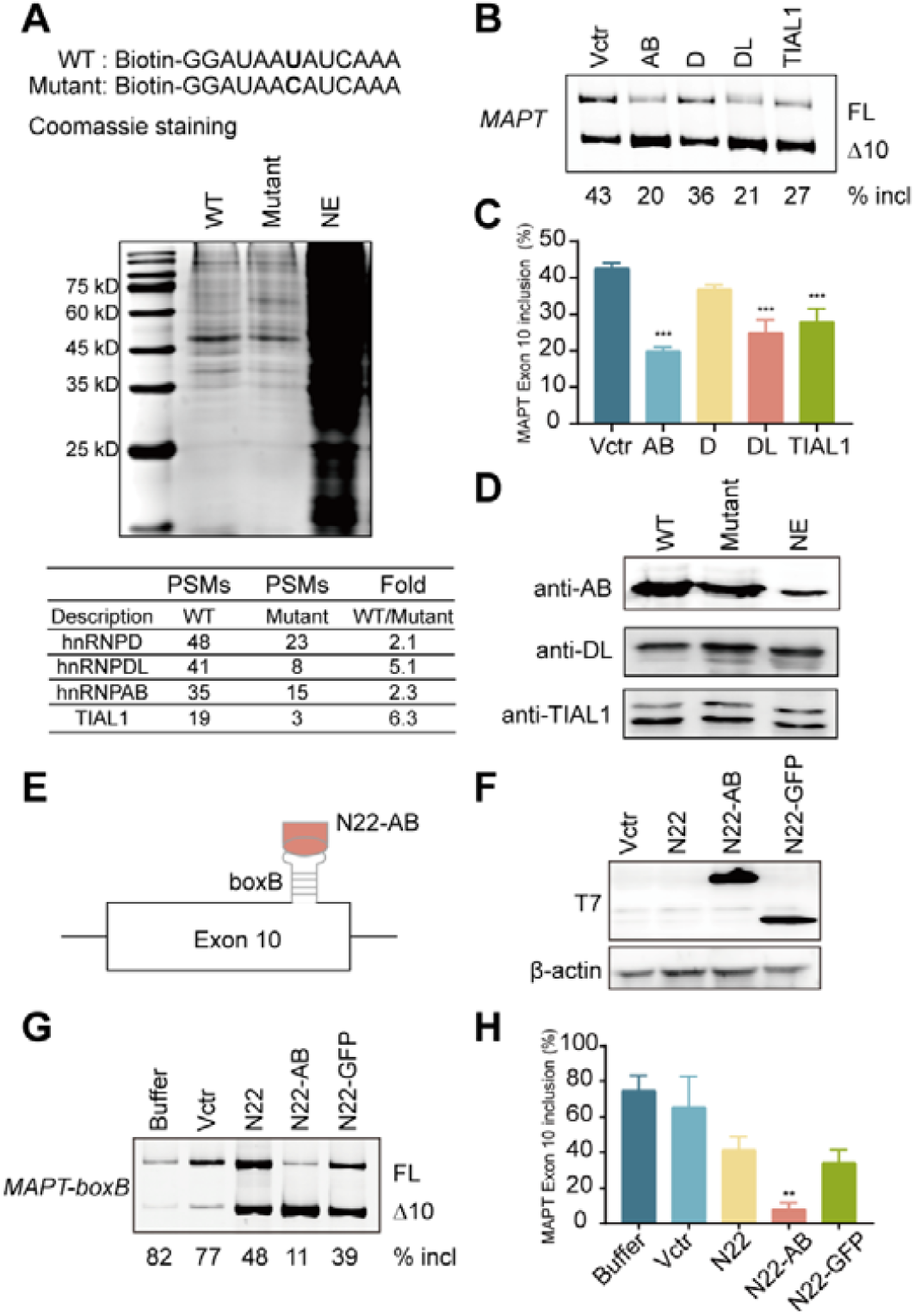
HnRNPAB binds to the AAUAU motif. **(A)** RNA pull-down assays were performed to identify the hnRNPAB-binding motif. **(B-C)** Analysis of *MAPT* exon 10 splicing regulation by the indicated expression plasmids. **(D)** Western blot analysis of eluted proteins confirmed specific enrichment of hnRNPAB in the WT RNA sample. **(E)** Schematic illustrating targeting of N22 fusion protein to *MAPT-boxB*. **(F)** Western blot analysis verified the proper expression of all fusion proteins in HeLa cells. **(G-H)** The effect of N22-fused proteins on the *MAPT*-*boxB* minigene was analyzed (n = 3). \*\**P* < 0.01, \*\*\**P* < 0.001 vs Vctr.

To verify that hnRNPAB exerts splicing repression through binding AAUAU motif, we employed a boxB-*λ*N protein tethering system as previously described (36,37). A mutant *MAPT* minigene (*MAPT-boxB*) was constructed by replacing exon 10 nucleotides +64 to +68 with a 19-nt boxB motif (Fig. 2E). HeLa cells were co-transfected with *MAPT-boxB* and T7-tagged N22-hnRNPAB or N22-GFP, and exon 10 splicing was analyzed. Fusion protein expression was confirmed by Western blot (Fig. 2F). This substitution significantly enhanced exon 10 inclusion efficiency, phenocopying the effect of ESS deletion. Although N22 displayed intrinsic repression of *MAPT-boxB* splicing, N22-hnRNPAB demonstrated potent suppression, reducing exon 10 inclusion from 42% (N22 control) to 8%, whereas T7-N22-GFP showed no regulatory activity (Fig. 2G-H). This confirms that N22-mediated tethering of hnRNPAB to the boxB motif specifically recapitulates WT *MAPT* minigene splicing repression, and it validates the mechanistic insights presented in Fig. 1.

### AB332 recruits RBMX/RBMXL1 via its RRM and Gly domains to repress *MAPT* exon 10 splicing

The *hnRNPAB* gene consists of 8 exons and encodes three isoforms—AB332 (the canonical isoform analyzed in Fig. 1, Fig. 2 and Fig. S1), AB327, and AB285—and the protein contains multiple domains, including a CARG-binding factor N-terminus domain (CBFNT), two central RNA recognition motifs (RRMs), and a C-terminal Gly-rich domain (Gly) (38). Relative to AB332, AB327 lacks 5 amino acids within the glycine rich domain. AB285, by contrast, lacks 47 amino acids within the glycine rich domain, a deletion caused by skipping of *hnRNPAB* exon 7 (Fig. S2A). To determine the functional relevance of these structural differences, we generated T7-tagged constructs for each isoform and evaluated their effects on endogenous and minigene *MAPT* splicing in HeLa cells. All isoforms repressed exon 10 inclusion in the *MAPT* minigene, but only AB332 strongly inhibited endogenous *MAPT* splicing (Fig. S2B-C). Moreover, AB332 overexpression retains its repressive activity even when all isoforms of hnRNPAB are knocked down, a capacity superior to that observed with AB285 overexpression. (Fig. S2D).

To identify AB332-specific binding proteins, we next performed immunoprecipitation (IP) (Fig. 3A). Seven proteins were enriched in the AB332 sample with PSMs ≥50 and AB332/AB285 PSM ratio >5: RBMX, RBMXL1, RPL4, TUFM, HADHB, EIF2S3 and EIF4A1 (Fig. 3B). Co-immunoprecipitation (Co-IP) demonstrated only a specific interaction between AB332 and RBMX/RBMXL1, while no binding activity was detected for AB285 (Fig. 3C and Fig. S3A-B). To define the molecular interaction between AB332 and RBMX, we conducted protein-protein docking simulations using PyMOL 2.5.3. The results pinpointed key hydrogen bonds, linking AB332 residues Y303, T107, and R248 to RBMX residues E3, R49, and S73, respectively. Furthermore, a stabilizing salt bridge was formed between AB332 R109 and RBMX D42 (Fig. 3D). A highly similar binding pattern emerged from analogous simulations between AB332 and RBMXL1(Fig. S3C). RBMX expression was downregulated upon AB332 overexpression (Fig. S3D-E). Overexpression of T7-tagged RBMX or RBMXL1 only weakly repressed *MAPT* exon 10 inclusion, and co-expression with AB332 did not further alter splicing compared to AB332 alone (Fig. S3F-G). These data suggest that AB332 acts as the dominant effector in regulating *MAPT* exon 10 splicing, while its strong repression masks potential contributions from RBMX/RBMXL1.

**Fig. 3.**
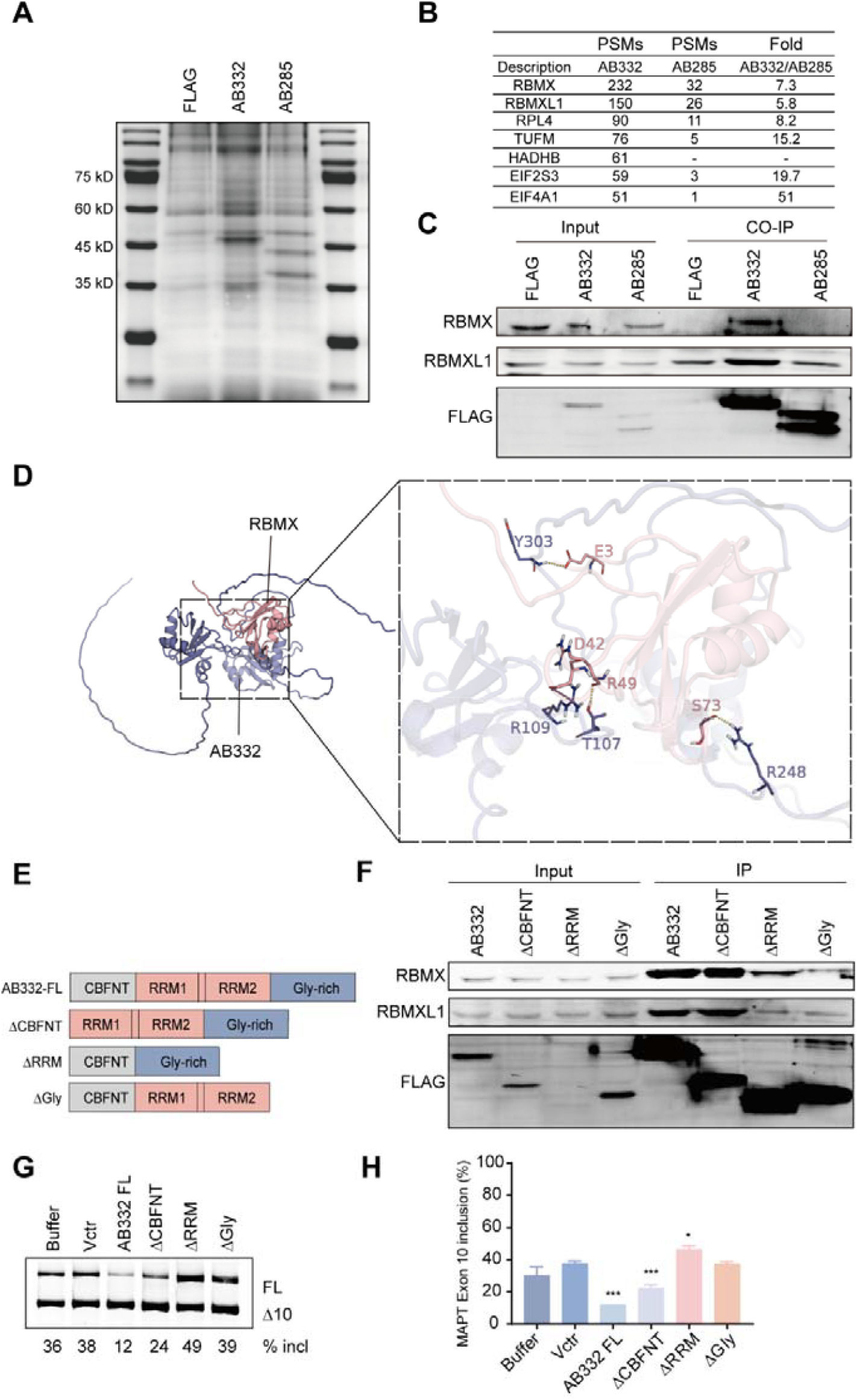
RRM and Gly domain of hnRNPAB bind RBMX/XL1 to inhibit *MAPT* exon 10 splicing. **(A-B)** IP assays were performed to identify hnRNPAB-interacting proteins. **(C)** Co-IP assays were performed using anti-RBMX and anti-RBMXL1 antibodies. **(D)** Molecular docking was used to simulate the binding mode between RBMX and AB332. **(E)** Schematic of the primary structure of hnRNPAB and its deletion mutants. **(F)** Co-IP assays were conducted to identify the key domains of AB332. **(G-H)** The effect of AB332 domain deletion mutants on *MAPT* minigene exon 10 splicing was analyzed in HeLa cells (n = 3). \**P* < 0.05, \*\*\**P* < 0.001 vs Vctr.

To map the interaction domain of the AB332 protein essential for RBMX/XL1 binding, we generated a series of FLAG-tagged deletion mutants targeting distinct structural regions (Fig. 3E). Co-IP showed that deleting the CBFNT domain of AB332 did not affect its binding to RBMX or RBMXL1. However, removing the RRM or Gly domain almost completely abolished these interactions (Fig. 3F). Consistently, the ΔCBFNT mutant repressed *MAPT* exon 10 splicing as effectively as AB332, whereas the ΔRRM and ΔGly mutants lost this repressive function (Fig. 3G–H). These results demonstrate that the RRM and Gly domains of AB332 mediate its functional interaction with RBMX/RBMXL1 to form an active splicing repressor complex.

### AB332 coordinates a network of alternative splicing events in AD-associated genes

To comprehensively characterize the regulatory effect of AB332, we performed isoform sequencing (Iso-seq) in HeLa cells stably overexpressing T7-tagged AB332. Relative to the Vctr, 484 genes were significantly downregulated and another 1438 were significantly upregulated (Fig. S4A). We used RT-qPCR to validate mRNA levels of *NT5E* and *NR1D1*, two genes implicated in AD pathogenesis (Fig. S4B-C). We performed differential transcript usage (DTU) analysis and identified 722 statistically significant altered splicing events involving 614 genes, with exon skipping (ES) accounting for 33.43%, intron retention (IR) for 12.75%, and other categories (including mutually exclusive exons (MXE) and alternative 5′/3′ splice sites (A5′SS, A3′SS)) for 53.82% (Fig. 4A). Among these altered events, 35.29% exhibited increased exon inclusion (ΔPSI > 0, *P* < 0.05), with the majority (64.71%) showing increased skipping (ΔPSI < 0, *P* < 0.05) (Fig. 4B-C). KEGG enrichment analysis revealed that these differentially alternatively spliced genes are involved in mitophagy, apoptosis, and other pathways (Fig. S4D). Notably, AB332 modulated splicing of several genes implicated in AD pathogenesis. These include: *ApoER2*, whose increased exon 19 inclusion is associated with improved cognition (39); *TMPO*, connected to cellular senescence (40,41); *zDHHC7*, a key regulator of synaptic plasticity (42); *MCL-1*, whose promotes mitophagy and mitigates A*β* pathology (43); *STAG2*, implicated in brain development (44); and the AD risk gene *PICALM*, whose loss impairs microglial clearance of A*β* and tau (45). Using RT-PCR, we validated that the three hnRNPAB isoforms regulate splicing changes, confirming that AB332 specifically increases inclusion of *ApoER2* exon 19 and *MCL-1* exon 2 and enhances skipping of specific exons in *TMPO, zDHHC7, STAG2*, and *PICALM*. In contrast, the AB285 isoform had no effect on these events (Fig. 4D-E). This consistent pattern confirms that the AB332 isoform specifically orchestrates splicing events relevant to AD pathways.

**Fig. 4.**
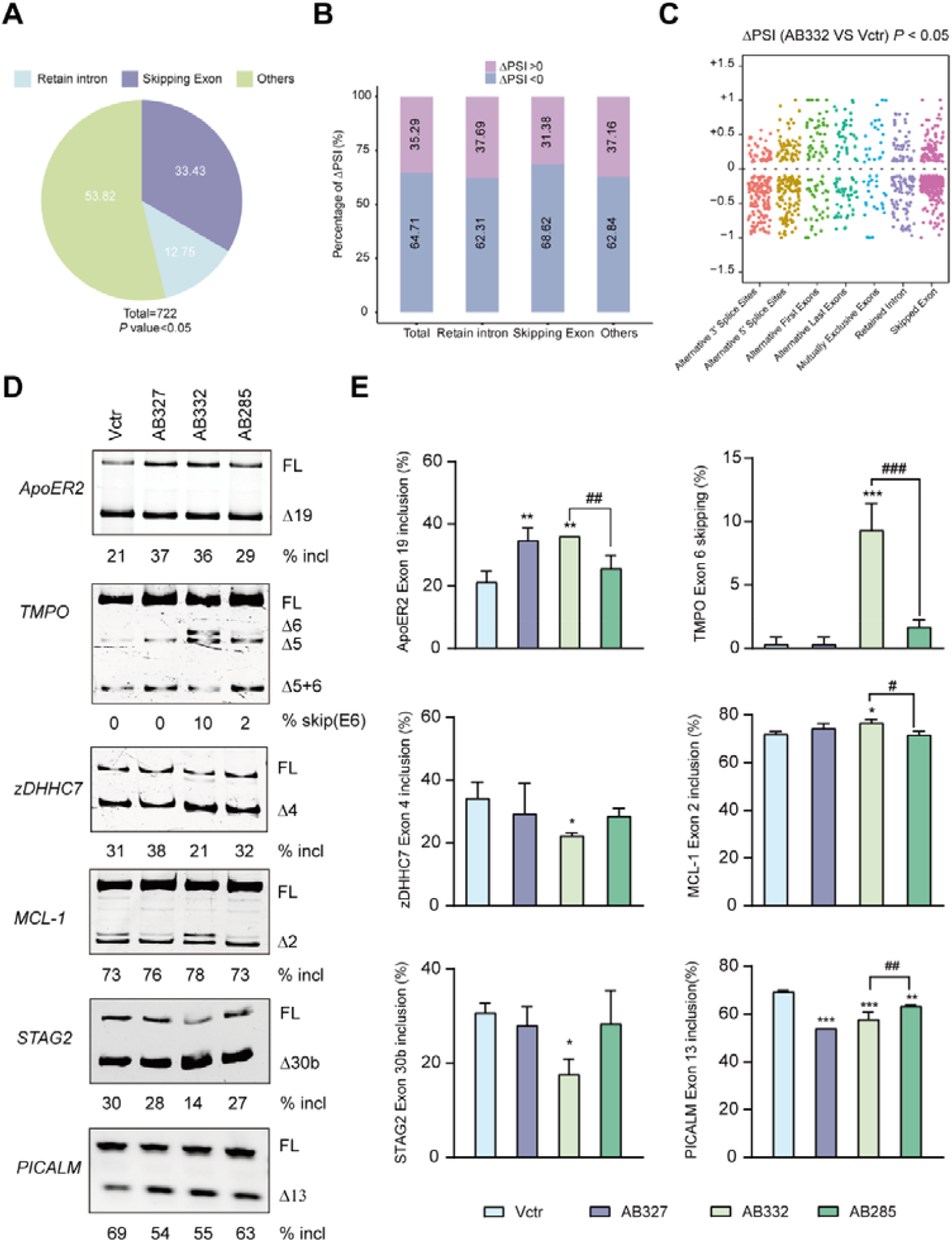
AB332 specifically regulates alternative splicing events of multiple AD-associated genes. **(A)** Pie chart showing the distribution of alternative splicing events (exon skipping, intron retention, alternative 5′/3′ splice sites, mutually exclusive exons) in HeLa cells overexpressing AB332. **(B)** Distribution of positive ΔPSI (PSI_AB332 OE_−PSI_Vctr_ > 0) and negative ΔPSI (PSI_AB332 OE_ − PSI_Vctr_ < 0) across total altered splicing events (n = 722), exon skipping (n = 241), intron retention (n = 92) and other splicing events (n = 389) (*P*-value< 0.05). **(C)** Distribution of ΔPSI values (PSI_AB332 OE_−PSI_Vctr_) for identified splicing events. Data are presented as median with interquartile range. **(D-E)** RT-PCR assays analyzing the regulation of alternative splicing events of *ApoER2, TMPO, zDHHC7, MCL-1, STAG2* and *PICALM* by different hnRNPAB isoforms in HeLa cells (n = 3 or 4). \**P* < 0.05, \*\**P* < 0.01, \*\*\**P* < 0.001 vs Vctr; #*P* < 0.05, ##*P* < 0.01, ### *P* < 0.001 vs AB332.

### Genome-wide mapping reveals AB332 binds AU-rich motifs to regulate splicing

To identify the direct RNA motif targets bound by AB332 isoform under physiological conditions, we performed enhanced crosslinking and immunoprecipitation sequencing (eCLIP-seq) in HeLa cells using an AB332-specific antibody. We identified 3022 genes with AB332 binding sites, predominantly enriched in the 3′ untranslated region (3′ UTR) near the stop codon (Fig. 5A-B). Peak annotation showed that about 33.78% of AB332 binding sites were located within exonic regions of coding sequences, and the vast majority of RNAs bound by AB332 were protein-coding transcripts (Fig. 5C-D). Motif analysis revealed a strong preference for AU-rich sequences, including the AAUAU motif identified in *MAPT* (Fig. 5E and Fig. S5A). We further intersected the eCLIP□seq targets with genes showing AB332□dependent alternative splicing changes from Iso-seq, identifying 165 overlapping genes, including *PICALM, TMPO, MCL-1*, and *STAG2* (Fig. 5F). GO and KEGG functional enrichment analyses of these 165 genes indicated AB332’s role in negatively regulating apoptosis (Fig. S5B-C). Finally, we visualized AB332 binding enrichment on *PICALM, TMPO, MCL-1*, and *STAG2* transcripts (Fig. 5G and Fig. S5D), providing direct evidence that AB332 binds their RNA to regulate splicing.

**Fig. 5.**
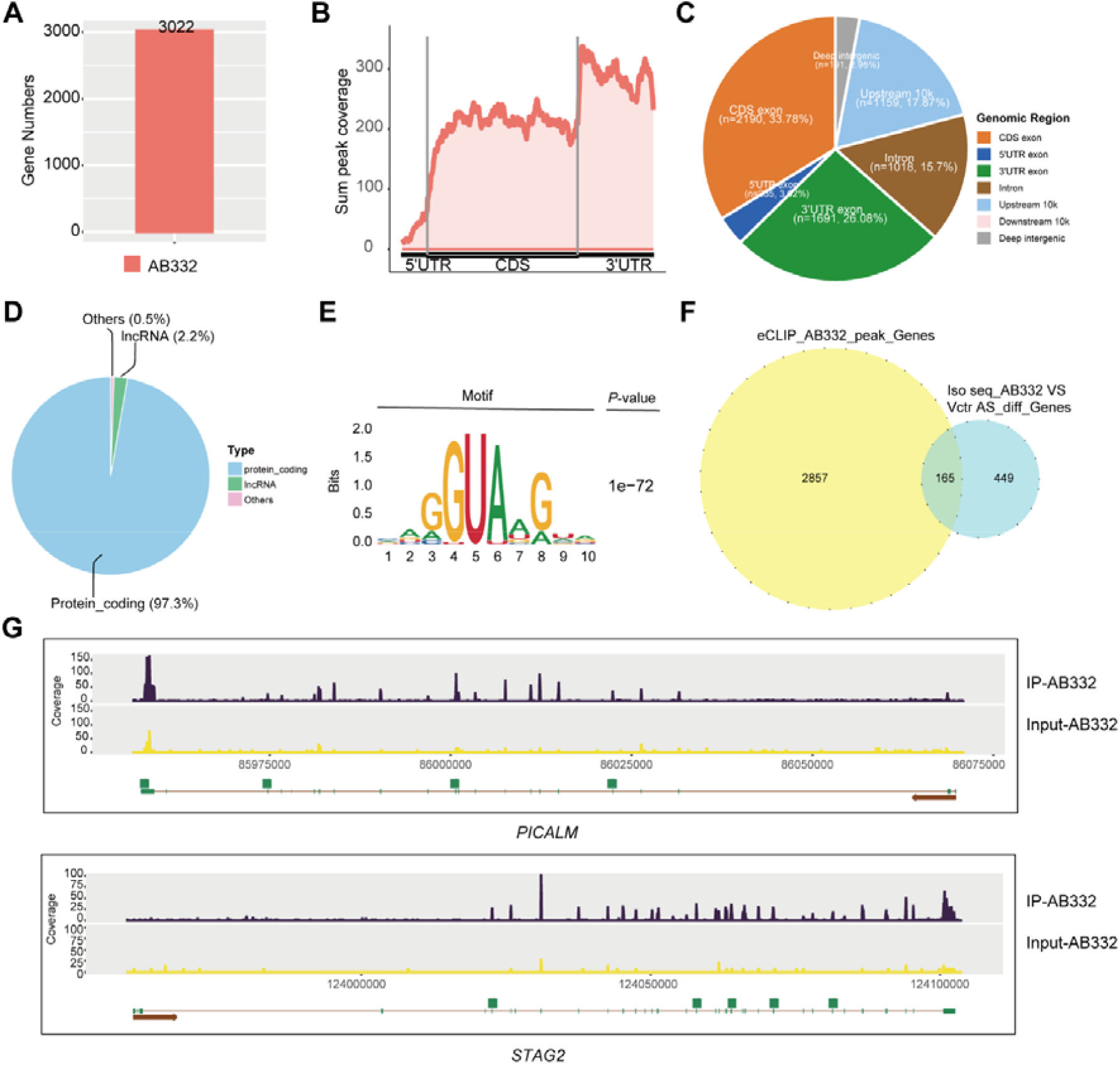
Identification of direct downstream targets of AB332 in HeLa cells. **(A)** Number of downstream genes harboring AB332-binding sites in HeLa cells. **(B)** Metagene profiles depicting the enrichment pattern of all AB332-binding peaks. **(C)** Pie chart illustrating the genomic distribution ratio of AB332-binding sites identified by eCLIP-seq. **(D)** Distribution of AB332 downstream target genes among the eCLIP-seq datasets. **(E**) Top significantly enriched AB332-binding motifs identified via de novo motif analysis in HeLa cells. **(F)** Overlapping genes between Iso-Seq and eCLIP-seq data for AB332 in HeLa cells. **(G)** eCLIP-seq read coverage of *PICALM* and *STAG2* genes in HeLa cells.

### Screening identifies ASO30 as a potent splicing modulator co-regulating *hnRNPAB* and *MAPT*

Given the central role of AB332 in mediating hnRNPAB-dependent splicing regulation, we explored its therapeutic potential in AD. We first analyzed the GEO transcriptomic dataset (GSE5281) of post-mortem brain tissues from AD patients, and found that *hnRNPAB* mRNA levels were significantly downregulated in AD samples compared to non-demented controls (Fig. S6A). We further detected the expression of the AB332 isoform in the hippocampus of 12-month-old female 3xTg-AD mice, with AB332 levels found to be selectively decreased when compared to age-matched wild-type littermates (Fig. S6B).

ASOs targeting *hnRNPAB* gene represent a promising AD therapeutic strategy by promoting exon 7 inclusion and upregulating AB332 expression. To implement this approach, we systematically screened a panel of ASOs targeting exon 7 and its flanking intronic sequences of the *hnRNPAB* gene in HeLa cells. This screen employed a three-step ASO walk method adapted from a previously reported protocol (46,47). We designed forty-two 22-mer overlapping ASOs, each modified with 2’-O-methoxyethyl ribose and a phosphorothioate backbone. These ASOs spanned a 266-nucleotide (nt) region, covering the final 62 nt of intron 6 through to the 63rd nt of intron 7. An unrelated oligonucleotide (ASO-00) served as a negative control (Fig. 6A and Fig. S7A). To assess ASO efficacy, we transfected HeLa cells with each ASO at 100 nM and analyzed *hnRNPAB* exon 7 splicing by semi-quantitative fluorescent RT-PCR. ASOs targeting exon 7 primarily induced skipping, whereas those directed against introns 6 and 7 generated more complex splicing patterns (Fig. 6B–C; Fig. S7B-C). Among the candidates, ASOs 29–32, 36, and 42 promoted *hnRNPAB* exon 7 inclusion to varying degrees. To identify the most effective candidate, we evaluated these six ASOs at 50 nM in both HeLa and SK-N-SH cells. At this concentration, only ASO30 significantly increased *hnRNPAB* exon 7 inclusion and promoted *MAPT* exon 10 skipping (Fig. 6D). ASO30 also showed cross-species activity, effectively modulating *hnrnpab* exon 7 splicing in murine BV2 and N2A cells despite a single-base mismatch (Fig. S8A-E).

**Fig. 6.**
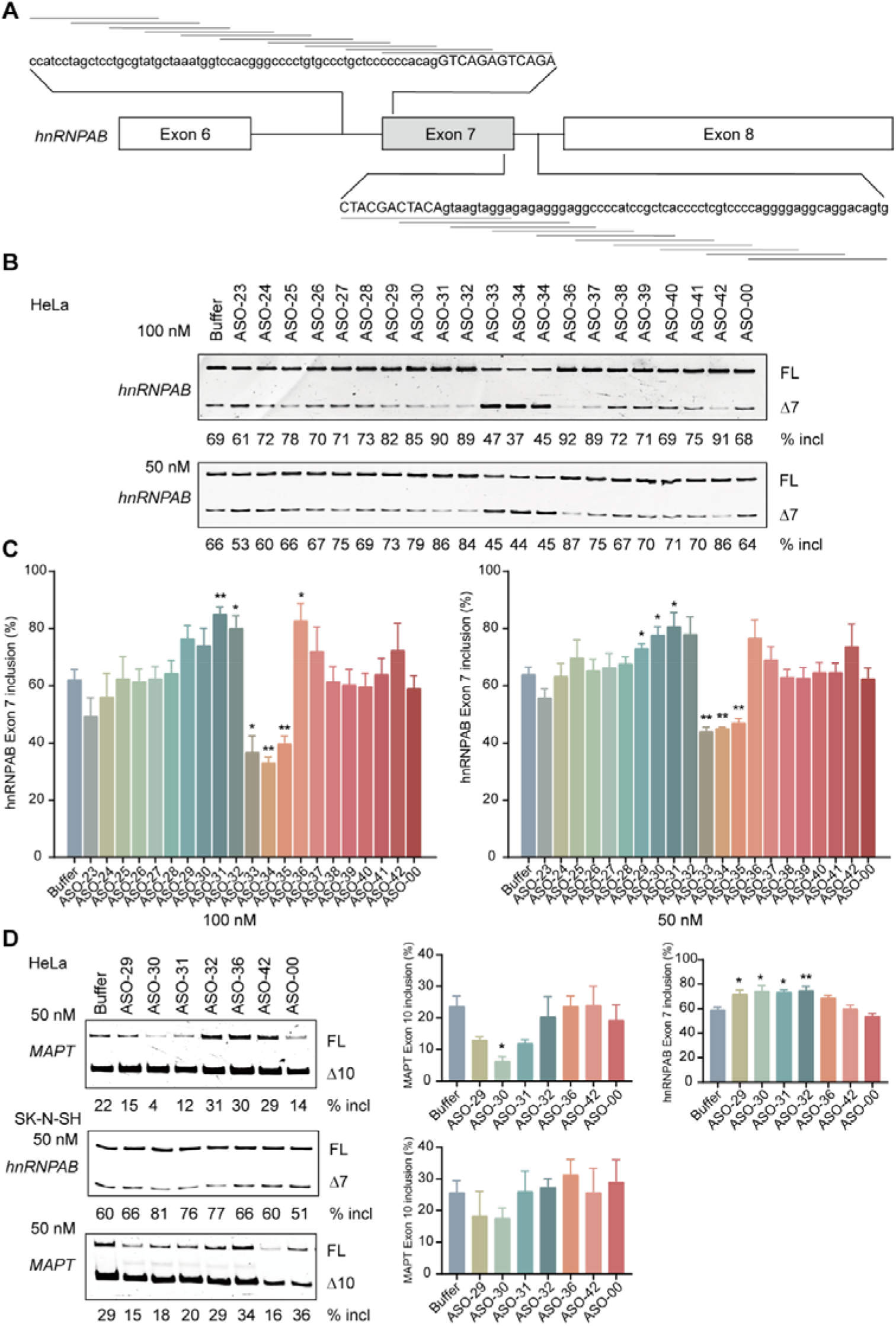
Systematic screening of ASOs that target *hnRNPAB* introns 6 and 7 and enhance *hnRNPAB* exon 7 inclusion. **(A)** Schematic of ASOs included in the screening. Each horizontal line corresponds to one individual ASO. **(B)** The impact of twenty individual ASOs at concentrations of 50 and 100 nM was examined in HeLa cells, with transfection buffer and an unrelated ASO (ASO-00) serving as negative controls. **(C)** Quantitative analysis of *hnRNPAB* exon 7 splicing efficiency corresponding to panel (B). **(D)** Splicing regulatory effects of six candidate ASOs (50 nM) on both *MAPT* and *hnRNPAB* transcripts in HeLa and SK-N-SH cells. (n = 3). \**P* < 0.05, \*\**P* < 0.01 vs buffer.

### ASO30 ameliorates cognitive deficits and tau pathology in 3xTg-AD mice

To determine the effective dose of ASO30 for in vivo studies, we performed a pilot dose-response experiment (0, 20, 50, 100, 200 ng) and found that 100 ng optimally increased *hnrnpab* exon 7 inclusion and promoted *mapt* exon 10 skipping (Fig. S8F-L). We then administered a single 100 ng dose of ASO30, a scrambled control oligonucleotide (ASO-NC), or saline via ICV injection to 9-month-old female 3xTg mice. After eight weeks, cognitive function in these mice was evaluated using a panel of behavioral tests (Fig. 7A). In the open field test, ASO30-treated mice spent more time in the central zone, indicating reduced anxiety-like behavior (Fig. 7B-C). The novel object recognition test revealed a restored preference for the novel object, demonstrating improved recognition memory (Fig. 7D-E). Spatial learning and memory, assessed by the Morris water maze, were also rescued: ASO30-treated mice crossed the former platform location more frequently, spent more time in the target quadrant (Fig. 7F-I). These behavioral improvements confirm that ASO30-mediated targeting of *hnrnpab* alleviates cognitive deficits in AD model.

**Fig. 7.**
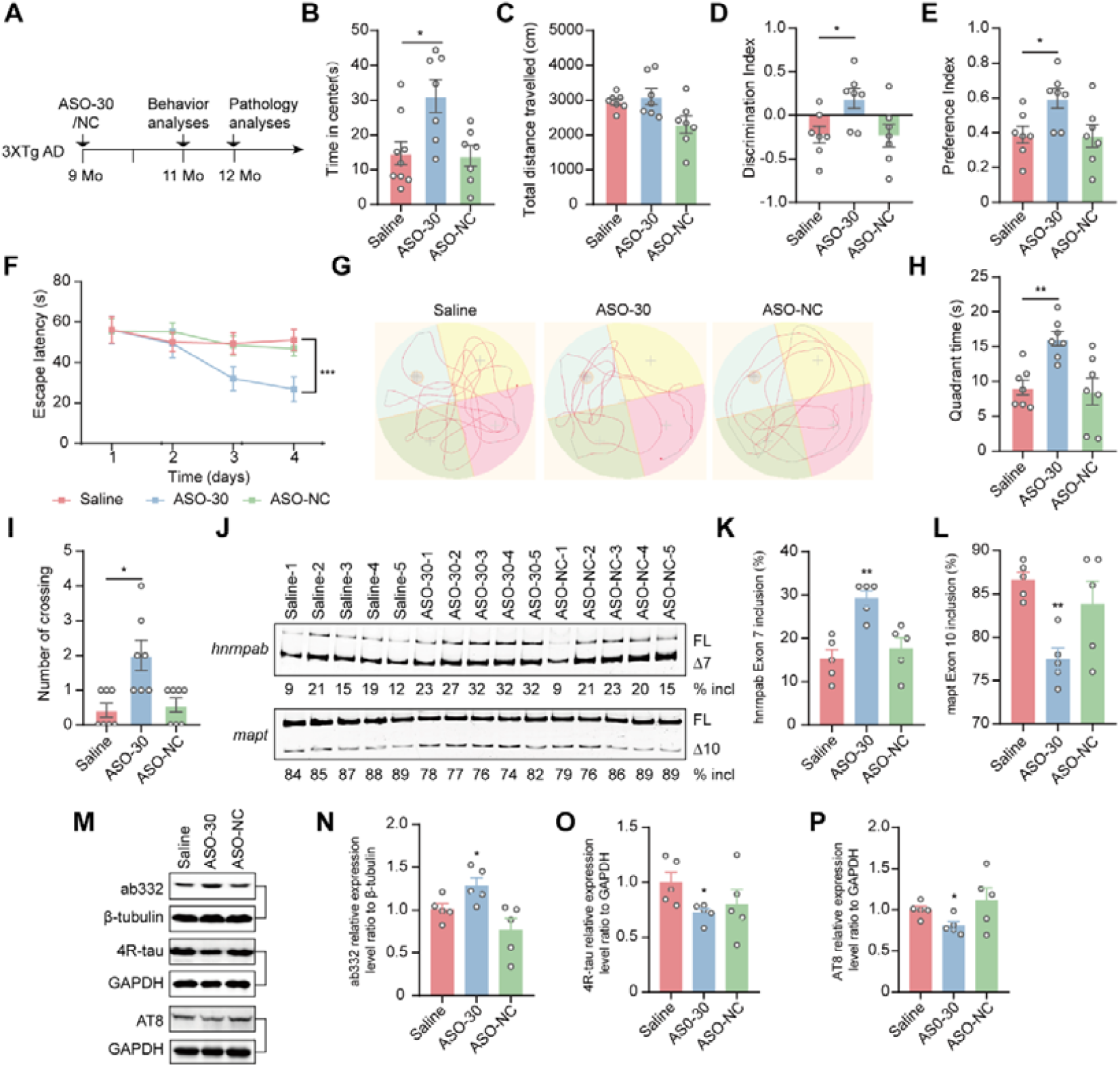
*hnrnpab*-targeting ASO ameliorates tau pathology and cognitive deficits in 3×Tg-AD mice. **(A)** Schematic diagram of unilateral ICV injection in 3×Tg-AD mice. **(B-C)** OF test to assess locomotor activity and anxiety-like behavior in 3×Tg-AD mice following ICV injection of ASO30 (n = 7). **(D-E)** NOR test to assess novel object preference in 3×Tg-AD mice following ICV injection of ASO30 (n = 7 mice per group). **(F-I)** MWM test to evaluate learning and memory abilities in 3×Tg-AD mice after ICV administration of ASO30 (n = 7). **(J-L)** RT-PCR analysis of the hippocampus to detect the splicing regulatory effects of ASO30 on *hnrnpab* and *mapt* genes in 3×Tg-AD mice following ICV injection (n = 5). **(M-P)** Westernblot analysis of ab332, 4R tau and p-tau proteins in the hippocampus of 3×Tg-AD mice receiving ICV injection of ASO30 (n = 5). \**P* < 0.05, \*\**P* < 0.01, \*\*\**P* < 0.001 vs Saline.

To link cognitive improvement to molecular mechanisms, we analyzed splicing and protein expression. ASO30 treatment significantly increased *hnrnpab* exon 7 inclusion and AB332 protein levels, promoted *mapt* exon 10 skipping, and reduced 4R-tau and phosphorylated tau levels (Fig. 7J-P). Collectively, these findings demonstrate that ASO30, through specific targeting of *hnrnpab*, effectively ameliorates both cognitive deficits and core tau pathology in AD.

## DISCUSSION

The hnRNP family comprises classical splicing regulators, several of which have been implicated in AD (48,49). However, the precise molecular mechanisms by which specific hnRNP members contribute to cognitive decline in AD remain to be fully elucidated. In this study, we establish hnRNPAB as a key regulator in this context. We demonstrate that its AB332 isoform forms a functional complex with RBMX/XL1 to recognize AAUAU motifs, thereby specifically repressing *MAPT* exon 10 inclusion and modulating a network of AD-associated splicing events. Correspondingly, we found that expression of AB332 isoform is significantly reduced in the hippocampus of AD model mice. Targeting this deficit with an antisense oligonucleotide (ASO30) designed to promote *hnRNPAB* exon 7 inclusion and upregulate AB332, we found that it effectively lowered phosphorylated tau levels and rescued cognitive deficits. These findings highlight the therapeutic potential of hnRNPAB-mediated correction of splicing dysregulation in AD.

Precise alternative splicing is orchestrated by the combinatorial action of splicing factors and SREs. Although regulation of *MAPT* exon 10 skipping—a critical event in tauopathy—has been extensively studied, with over 30 regulatory proteins reported, our work adds a new layer of complexity (50). We identified AB332 as a strong repressor of *MAPT* exon 10 inclusion, yet our previous mass spectrometry analyses established its well-characterized role as a potent activator of *SMN2* exon 7 splicing (36). This functional switch from activator to repressor is most likely attributable to a position-dependent effect, a well-established mechanistic principle wherein the splicing outcome dictated by an RNA binding protein is determined by its binding site relative to the regulated exon (51). Our findings align with this model and are consistent with prior examples such as TDP-43 and NOVA (52,53). Computational analyses, including those using tools like MAPP, have formally characterized this mechanism, showing how RNA binding proteins like RBFOX1 can exert opposite effects based on binding site context (54). Thus, AB332 exemplifies a key functional property of splicing regulators: context-dependent dual functionality, which allows a limited number of proteins to govern a vast array of splicing outcomes.

To established the context-dependent functionality of AB332, we next defined the molecular mechanism underlying its splicing repression of disease-associated exons. Deletion mapping pinpointed a critical AAUAU motif localized 64–68 nucleotides downstream of *MAPT* exon 10 as the key element of AB332 responsiveness. This finding is strongly supported by prior work. Lisowiec et al. reported that this element contributes to the formation of H4.2—a repressive hairpin structure with exon splicing silencer activity—whose upstream sequence recruits splicing factors including SRSF6, RBMX or/and SRSF9 (55); Wang et al. also identified this position as an exon splicing silencer through the MAPT N296N(T-C) mutation (56), which aligns with our finding that the thymine at position 66 of MAPT exon 10 plays an essential role within this regulatory element (Fig. 1). Our data extend the understanding of hnRNPAB binding specificity, as the AAUAU core we identified is embedded within the broader consensus sequence (SCCGMM) characterized by Fukuda et al. in the 3′UTRs of mRNAs associated with neural development (33). This AAUAU motif was also recovered in our independent eCLIP-seq analysis, confirming its status as a valid AB332 binding site. While the full spectrum of its binding motifs requires further study, our work definitively establishes this AAUAU element as a functional motif mediating hnRNPAB-dependent repression of MAPT exon 10 splicing.

RBMX is well-established to regulate PKM splicing through competitive binding with hnRNPA1 (57). Our present work further reveals that RBMX assembles into a functional complex with AB332 to repress MAPT exon 10 inclusion. We observed that AB332 overexpression suppresses RBMX expression (Fig. S3), consistent with a negative feedback regulatory loop. Of note, RBMX expression is reportedly increased in AD (58). We propose that downregulation of AB332 in the disease state may alleviate this inhibitory effect, leading to compensatory upregulation of RBMX. This dysregulated expression profile ultimately compromises the functional integrity of the AB332–RBMX complex, contributing to aberrant MAPT splicing in AD. We demonstrated that the interaction between AB332 and RBMX depends critically on the RRM and Gly domains of AB332. This finding aligns with a recent study by Pan et al., which established that the RRM1 domain of hnRNPAB is critical for its interaction with TRIM25 (30). It is also consistent with work from Wang et al. that identified the Gly domain of hnRNPAB as a key region mediating influenza A virus replication inhibition (38).

Our Iso-seq analysis identified a specific regulatory effect of AB332, which modulates the expression of AD-associated genes—an activity absent in the AB285 isoform. We focused particularly on two target genes: NR1D1 and NT5E. Overexpression of AB332 in HeLa cells elevated NR1D1 levels. This gene encodes REV-ERBα, a protein with complex, context-dependent roles in AD that involve processes such as autophagy and tau clearance (59-62). The effect of AB332 to influence NR1D1 suggests it may participate in these pathways, though its precise role across different cell types and disease stages requires further study. Similarly, AB332 upregulated NT5E, a gene whose reduced expression is known to exacerbate AD pathology (63). This consistency between our data and established pathological mechanisms strengthens the relevance of AB332 in AD. Furthermore, AB332 modulates alternative splicing of multiple AD-associated genes, including ApoER2, zDHHC7, MCL-1, and STAG2. Among these targets, MCL-1 is known to regulate both pathways for alleviating AD-related cognitive impairment—mitophagy and apoptosis—wherein its alternative splicing generates the anti-apoptotic MCL-1L and a shorter pro-apoptotic isoform, and it functions as a mitophagy receptor (64-68). Our study demonstrates that AB332 may directly bind MCL-1 mRNA and promote its splicing toward the protective MCL-1L isoform (Fig. 4 and Fig.5), thereby likely suppressing apoptosis and enhancing mitophagy, and our Iso-seq further supports this view by showing that genes with AB332-altered splicing are enriched in autophagy and mitophagy pathways (Fig. S4). Our results suggest that AB332 functions as a key splicing regulator that connects to established neuroprotective mechanisms, although the precise molecular basis of this regulatory crosstalk remains to be fully elucidated.

Previous clinical successes demonstrate the promise of ASO therapy for neurological diseases. Nusinersen, a 2′-MOE/PS-modified ASO, corrects SMN2 splicing and is an approved treatment for spinal muscular atrophy (47,69). Similarly, the ASO MAPTRx reduces total tau levels in AD patients (70). However, non-selectively lowering tau may disrupt its essential physiological roles in neuronal structure and function (71,72). A safer strategy would selectively reduce pathological tau while sparing its normal function. In this study, the hnRNPAB-targeting ASO embodies such an approach by modulating the alternative splicing of multiple AD-associated genes, including by promoting MAPT exon 10 skipping, reducing 4R-tau and phosphorylated tau levels, and ultimately enhancing cognitive function. This strategy may thus be broadly applicable to other tauopathies, including frontotemporal dementia.

In summary, we identified a candidate ASO targeting hnRNPAB alternative splicing as a potential therapeutic strategy for AD and related tauopathies. The main limitation of this study is that we did not assess the duration of ASO action or further optimize the candidate ASO in 3xTg mice, as this study focused on identifying hnRNPAB as a novel therapeutic target for AD. In the future, this approach may be further optimized by combining the ASO with targeted delivery strategies—for instance, low-intensity focused ultrasound. Such a combined strategy could improve drug efficacy by enhancing ASO bioavailability in the brain, while simultaneously enhancing patient compliance and safety.

## ACKNOWLEDGEMENTS

The author declares no additional acknowledgments for this work.

## AUTHOR CONTRIBUTIONS

Tao Jiang: Designed the study, Performed experiments, Data analysis, Writing. Xingjun Xu: Performed experiments, Data analysis. Yaqiong Yu: Performed experiments. Caixin Zhu: Data analysis. Chao Ma: Data analysis. Yu Tang: Data analysis. Lingxia Min: Data analysis. Mingliang Tan: Data analysis. Jialin Bai: Writing. Zhou Feng: Designed the study, Writing. Jingming Hou: Designed the study, Writing.

## SUPPLEMENTARY DATA

Supplementary Data are available at NAR online.

## DATA AVAILABILITY

All data associated with this study are present in the paper or the Supplementary Materials. Sequencing data are deposited at SRA: Iso-seq data accession number PRJNA1423612; eCLIP-seq data accession number PRJNA1423657.

