## Supplemental Figures and Tables for "*HnRNPAB*-targeted antisense oligonucleotides ameliorate tau pathology and cognitive deficits by modulating Alzheimer’s disease-associated alternative splicing"

**This PDF file includes:**

Figures S1 to S8
Tables S1 to S4

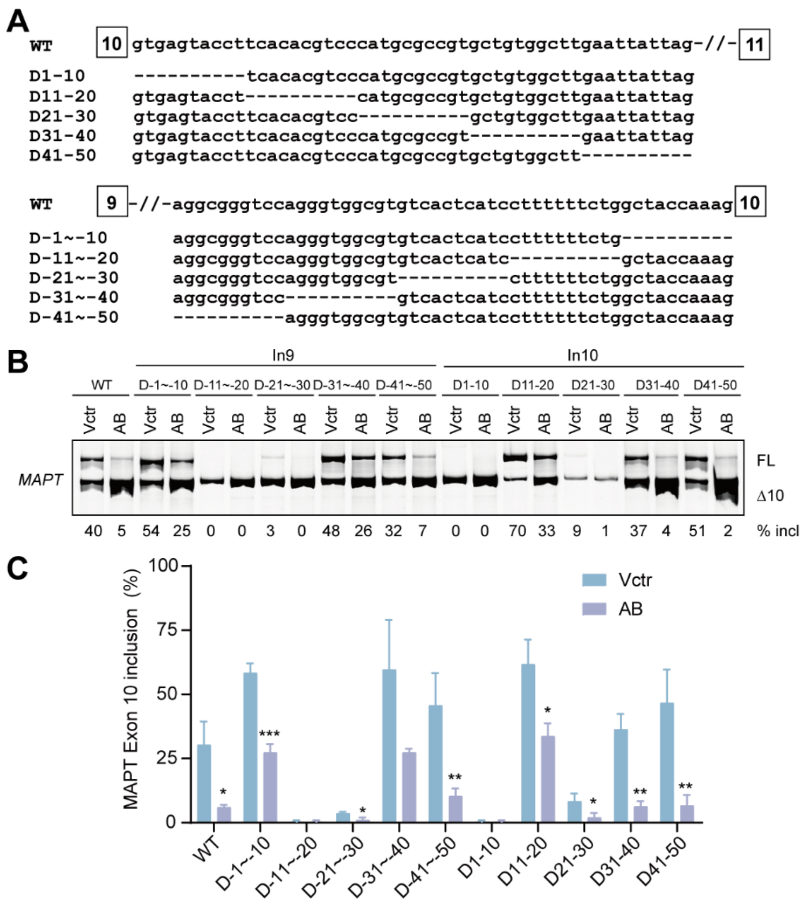


**Fig. S1. Mutational analysis of regions spanning intron 9 and intron 10 of the *MAPT* minigene.** (**A**) Sequences of the intron 9 and intron 10 deletion mutants. (**B**) HeLa cells were co-transfected with each mutant or wild-type (WT) minigene plasmid plus the T7-tagged hnRNPAB expression plasmid; empty vector (Vctr) served as the negative control. *MAPT* exon 10 splicing patterns were analyzed via Cy5-labeled RT-PCR. (**C**) Histogram presents quantitative data from three independent experiments corresponding to panel (B) (n = 3). *$P$ < 0.05, **$P$ < 0.01, ***$P$ < 0.001. FL: full length; Δ10: exon 10-skipped isoform; % incl: percentage of exon 10 inclusion; AB: T7-tagged hnRNPAB.

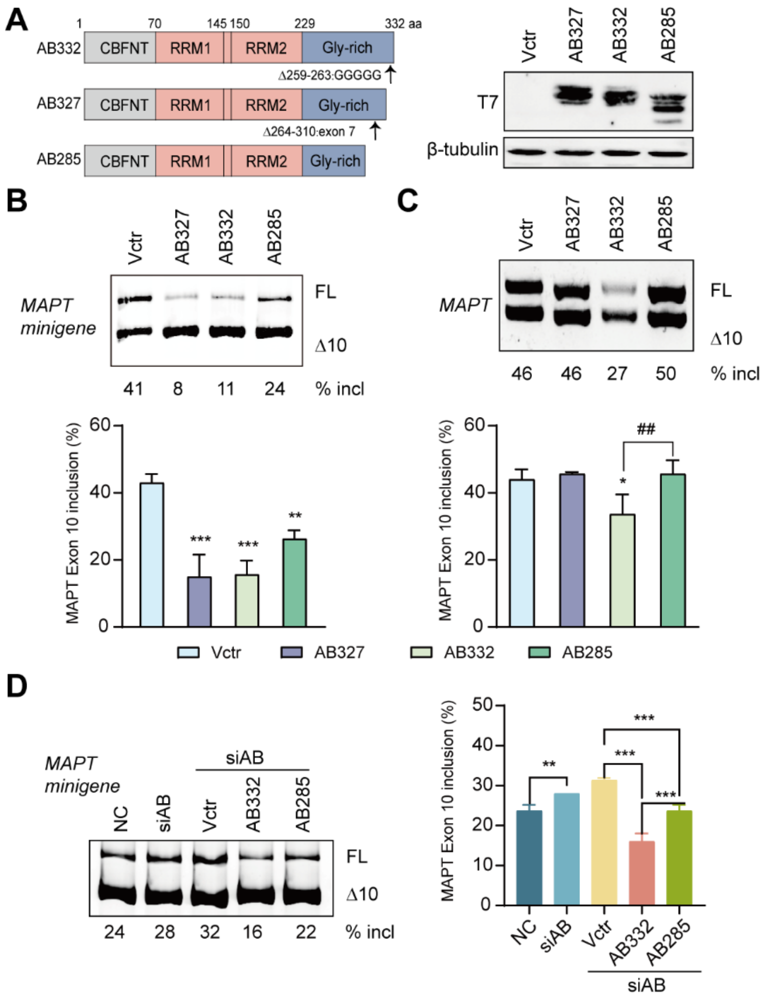


**Fig. S2. Analysis of the effects of hnRNPAB isoforms on *MAPT* splicing.** (**A**) Diagram of the primary structure of hnRNPAB and its three splicing isoforms. (**B**) Effects of overexpressing three hnRNPAB isoforms on *MAPT* minigene splicing in HeLa cells. (**C**) Effects of overexpressing three hnRNPAB isoforms on endogenous *MAPT* splicing in HeLa cells. Quantitative analysis of splicing data for hnRNPAB isoforms (n = 3). *$P$ < 0.05, **$P$ < 0.01, ***$P$ < 0.001 vs Vctr. ## $P$ < 0.01 vs AB332. (**D**) Splicing patterns of *MAPT* minigene in HeLa cells first treated with target-specific siRNA to deplete endogenous hnRNPAB, then transfected with T7-tagged hnRNPAB isoform expression plasmids or Vctr in the same culture. Non-targeting siRNA served as the negative control; target siRNA plus empty vector was used as the knockdown-only control. (n = 3). **$P$ < 0.01, ***$P$ < 0.001.

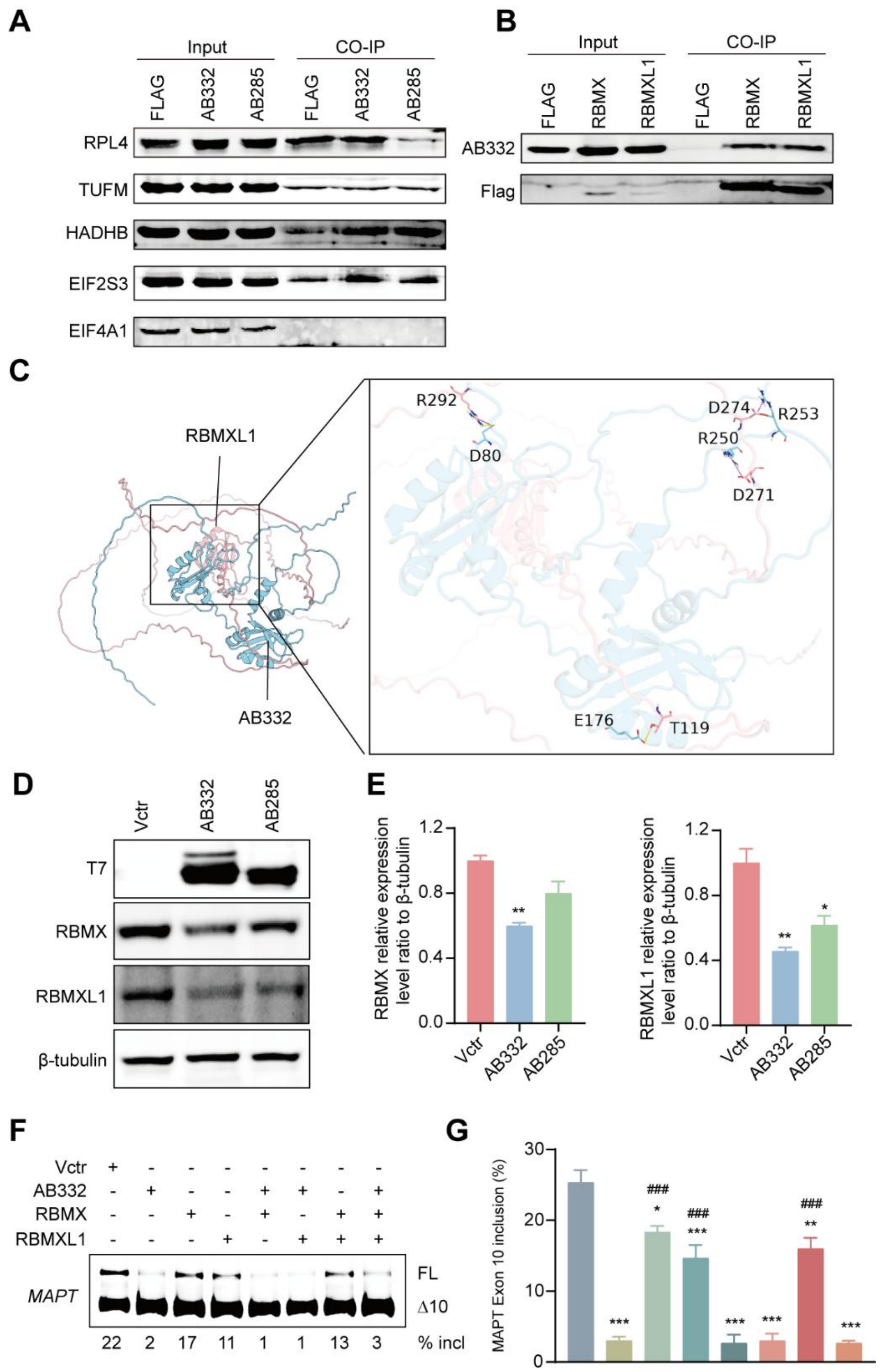


**Fig. S3. Identification of proteins interacting with AB332.** (**A**) Co-IP assays were performed to verify the interaction between RPL4, TUFM, HADHB, EIF2S3, EIF4A1 and AB332. (**B**) FLAG-RBMX and FLAG-RBMXL1 were used in Co-IP assays to capture AB332, validating the interaction of RBMX/XL1 with AB332. (**C**) Molecular docking was used to simulate the binding mode between RBMXL1 and AB332. (**D**) HeLa cells were transfected with T7-tagged AB332, T7-tagged AB285 or T7-tagged empty vector (as the negative control). Western blot assays were performed to detect the expression levels of RBMX and RBMXL1. (**E**) Quantitative analysis of RBMX and RBMXL1 expression levels corresponding to panel (D) (n = 3). (**F**) Effect of overexpressing T7-tagged AB332, RBMX or RBMXL1 alone (300 ng of each plasmid transfected), or their pairwise and triple combinations (150 ng or 100 ng of each plasmid co-transfected per group) on *MAPT* minigene splicing in HeLa cells, with T7-tagged empty vector (300 ng) as the negative control. (**G**) Quantitative analysis of *MAPT* exon 10 splicing efficiency corresponding to panel (F) (n = 3). *$P < 0.05$, **$P < 0.01$, ***$P < 0.001$ vs Vctr; ### $P < 0.001$ vs AB332.

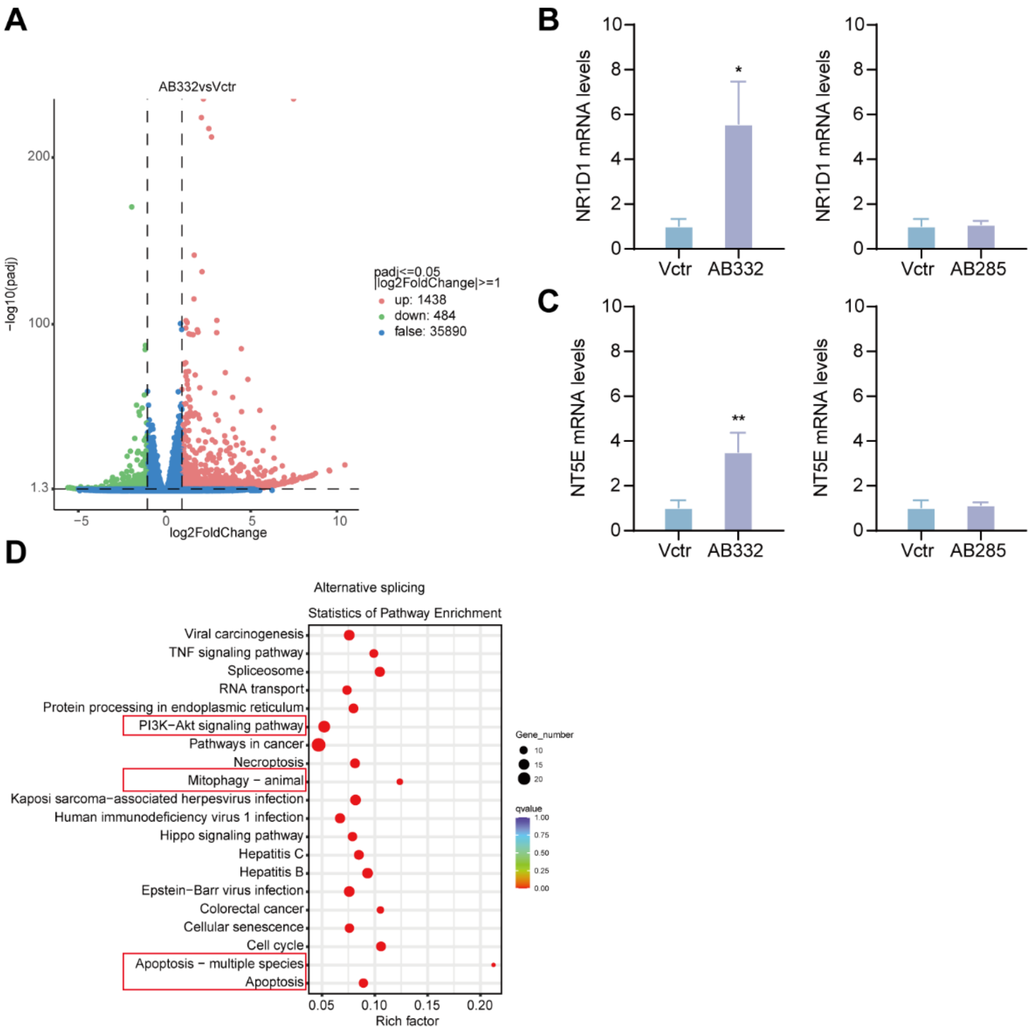


**Fig. S4. Identification of DEGs and dysregulated alternative splicing events in AB332-overexpressing**. (**A**) Volcano plot of differentially expressed genes (DEGs; |log2FoldChange|>=1, adjusted $P$ < 0.05) identified by RNA-seq analysis of Vctr and AB332 overexpression groups. Green dots indicate downregulated genes, and pink dots represent upregulated genes. (**B-C**) Quantitative real-time PCR (qRT-PCR) validation of selected DEGs. (n = 3). *$P$ < 0.05, **$P$ < 0.01 vs. Vctr. (**D**) KEGG enrichment terms for genes with dysregulated alternative splicing events in the AB332 overexpression group.

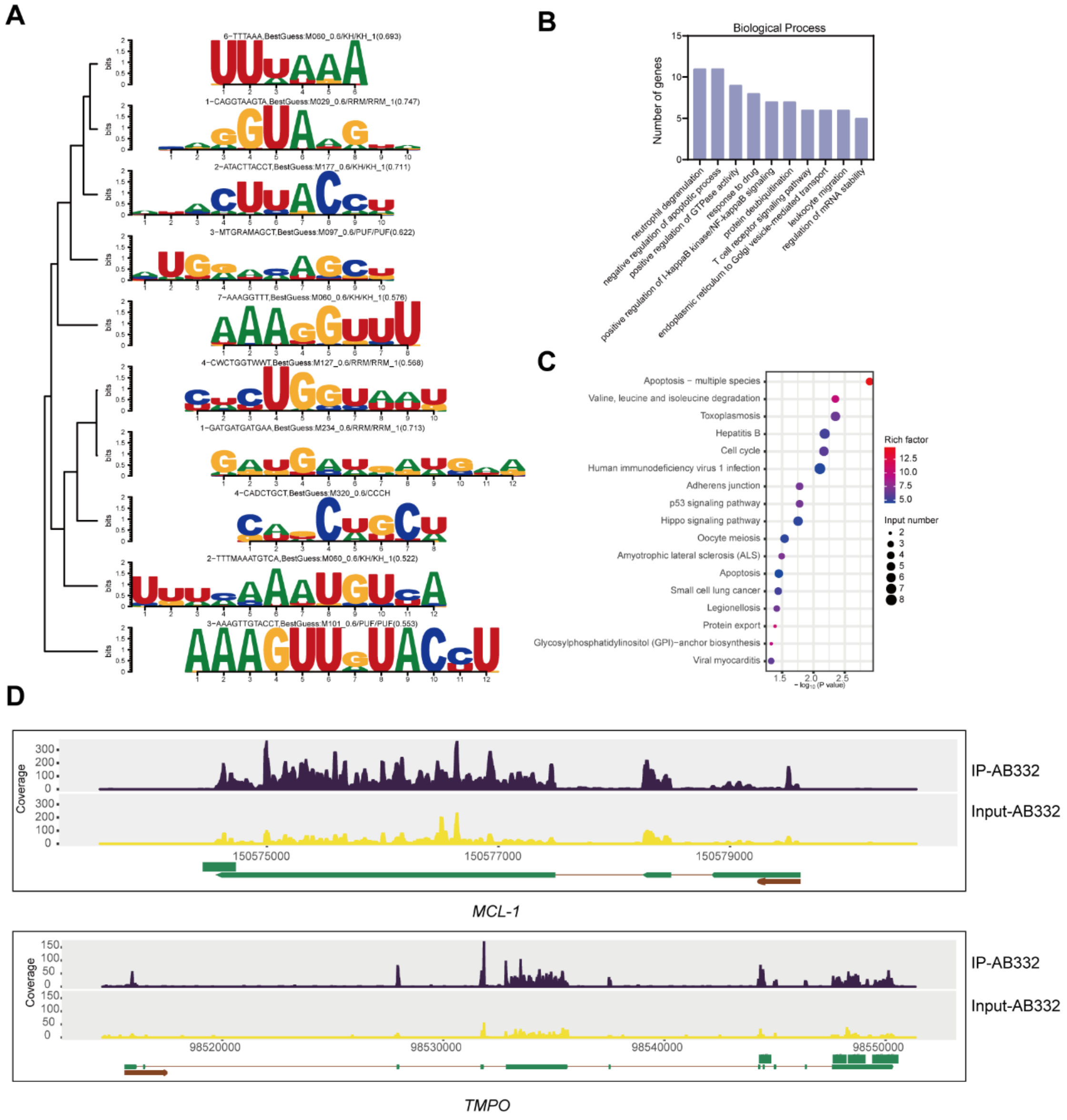


**Fig. S5. De novo motif discovery and functional enrichment analysis of AB332-binding target genes.** (**A**) The top 10 motifs identified de novo from AB332 eCLIP-seq peaks via HOMER analysis. (**B**) Top enriched pathways of genes harboring the AB332 binding consensus motif, as identified by GO biological process enrichment analysis. (**C**) Top enriched pathways of genes harboring the AB332 binding consensus motif, as identified by KEGG pathway enrichment analysis. (**D**) eCLIP-seq read coverage of the genomic regions of *MCL-1* and *TMPO* genes in HeLa cells.

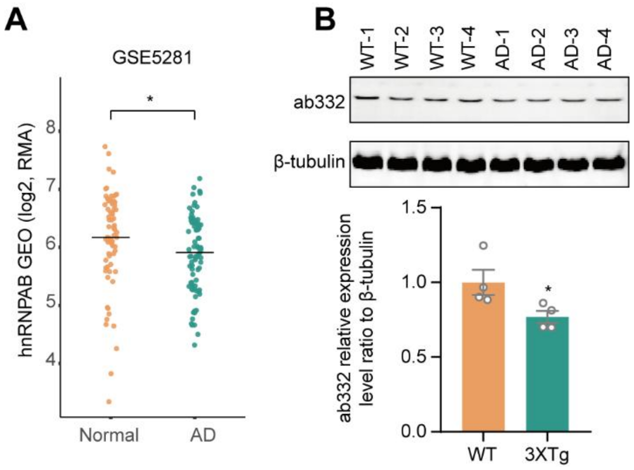


**Fig. S6. Expression of AB332 is decreased in AD.** (**A**) Transcription levels of *hnRNPAB* in AD patients were evaluated using GEO datasets. (**B**) Western blot analysis of AB332 protein expression in the hippocampus of WT and 3×Tg-AD mice (n = 4). Quantitative analysis of AB332 expression levels. *$P$ < 0.05 vs WT.

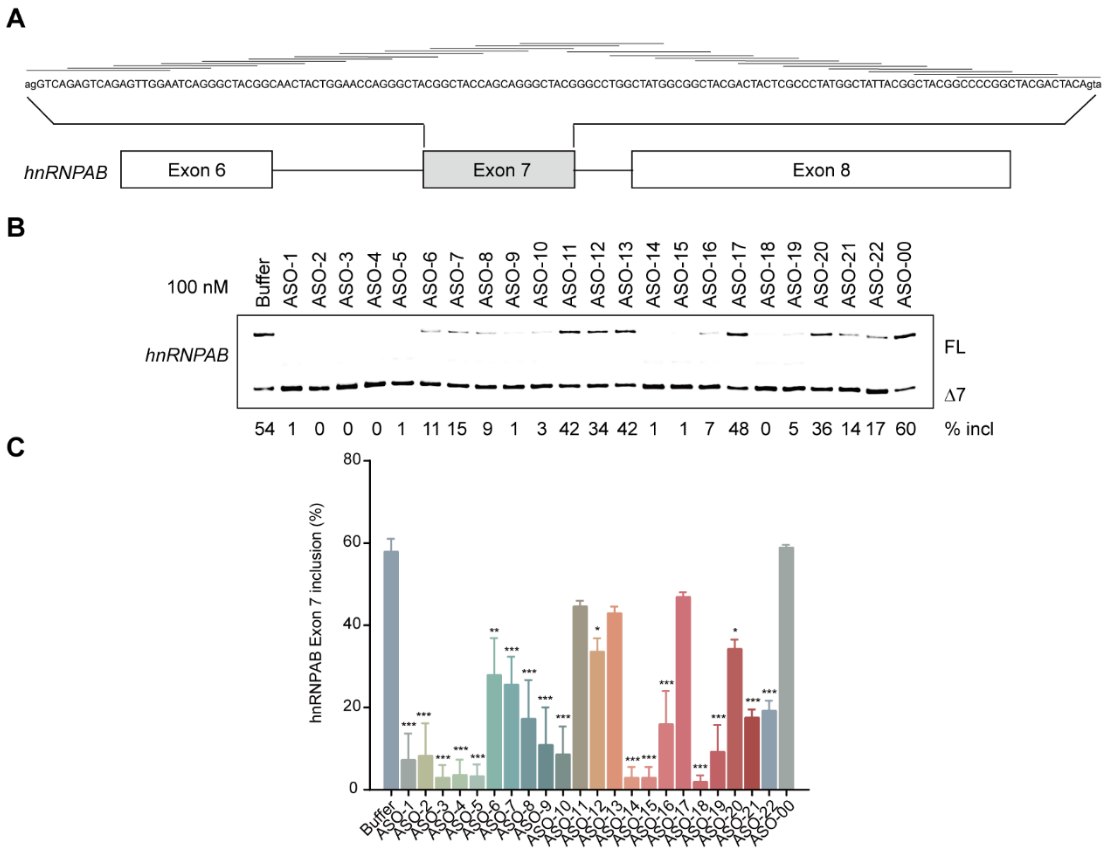


**Fig. S7. Systematic screening of ASOs targeting *hnRNPAB* exon 7.** (**A**) Schematic representation of ASOs used in the initial screening assay. Each horizontal line represents a single ASO sequence. (**B**) The effect of twenty-two ASOs at a concentration of 100 nM was evaluated in HeLa cells, with transfection buffer and an unrelated ASO (ASO-00) included as negative controls. (**C**) Quantitative analysis of *hnRNPAB* exon 7 splicing efficiency corresponding to panel (B) (n = 3). *$P < 0.05$, **$P < 0.01$, *** $P < 0.001$ vs buffer.

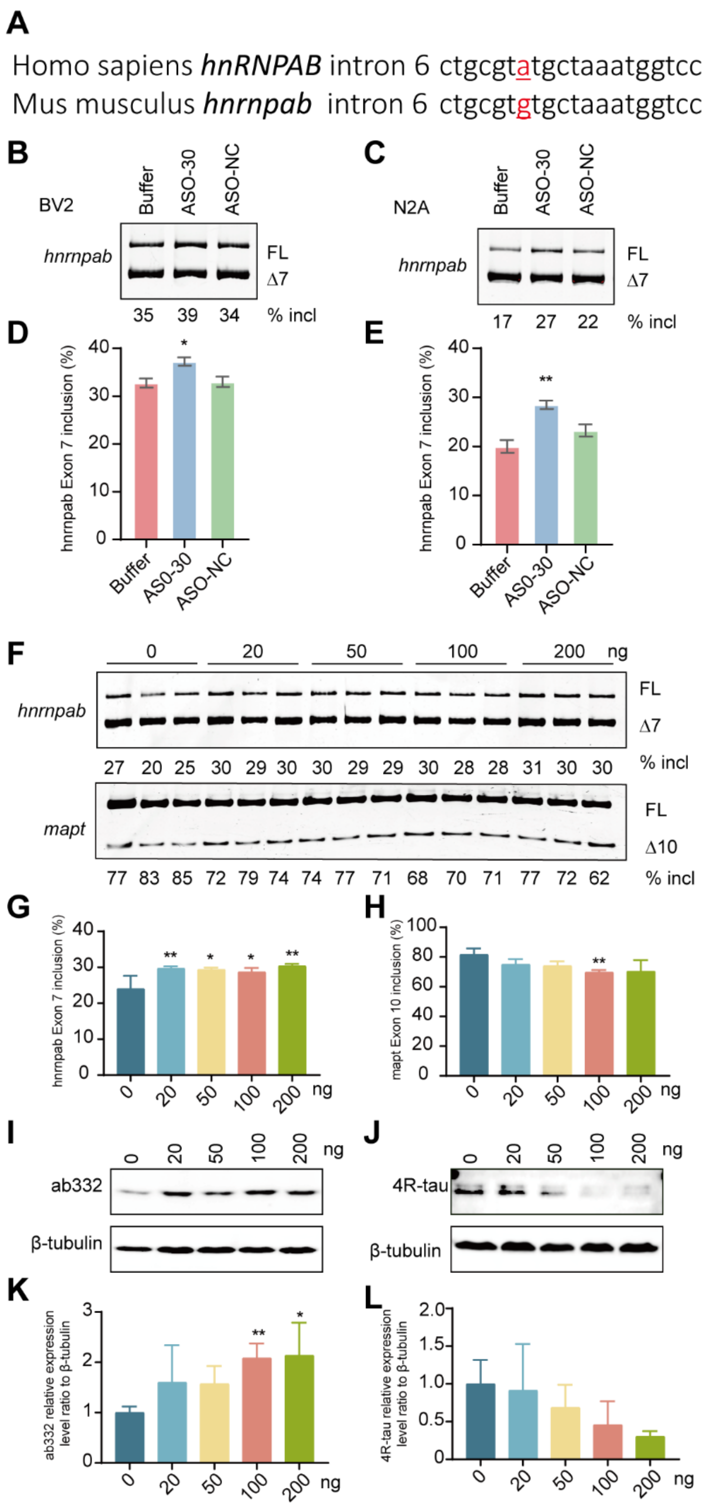


**Fig. S8. ASO-30 mediates *hnrnpab/mapt* splicing in murine cells and ICV dose effects in mice.** (**A**) Sequence alignment of *hnRNPAB* intron 6 targeted by ASO30**.** (**B**) The effects of ASO30 at a concentration of 50 nM in BV2 cells. (C) The effects of ASO30 at a concentration of 50 nM in N2A cells. (**D**) Quantitative analysis of *hnrnpab* exon 7 splicing efficiency corresponding to panel (B). (n=4 ). $^{*}P < 0.05$ vs buffer. (**E**)

Quantitative analysis of *hnrnpab* exon 7 splicing efficiency corresponding to panel (C). (n=4 ). **$P < 0.01$ vs buffer. (**F-H**) RT-PCR analysis of alternative splicing changes of *hnrnpab* and *mapt* in the hippocampus at 6 weeks post-injection across ASO-30 doses of 0, 20, 50, 100 and 200 ng per mouse. (n = 3). *$P < 0.05$, **$P < 0.01$ vs 0 ng control group. (**I-L**) Western blot analysis of expression changes of ab332 and 4R-tau proteins following treatment with the same ASO-30 dose gradient (n = 3). *$P < 0.05$, **$P < 0.01$ vs 0 ng control group.

**Table S1: Primer sequences used for plasmids construction.**

| Name | Sequence (5′ to 3′) |
|---|---|
| HnRNPAB-F | ATGTCGGAAGCGGGCGAG |
| HnRNPAB-R | TCAGTATGGCTTGTAGTTATTCTGATGG |
| SLIC T7 HNRNPAB-F | TGACAGGTGGCCAACAGATGGGTATGTCGGAAGCGGGCGAG |
| SLIC T7 HNRNPAB-R | GAGGGTCCATGGTGATACAAGGGATCAGTATGGCTTGTAGTTATTCTGATGG |
| SLIC T7-F | tcccttgtatcaccatggaccctc |
| SLIC T7-R | CATCTGTTGGCCACCTGTCA |
| HnRNPD-F | ATGTCGGAGGAGCAGTTC |
| HnRNPD-R | TTAGTATGGTTTGTAGCTATTTTG |
| SLIC T7 HnRNPD-F | TGACAGGTGGCCAACAGATGGGTATGTCGGAGGAGCAGTTC |
| SLIC T7 HnRNPD-R | GAGGGTCCATGGTGATACAAGGGATTAGTATGGTTTGTAGCTATTTTG |
| HnRNPDL-F | ATGGAGGTCCCGCCCAGG |
| HnRNPDL-R | TTAGTATGGCTGGTAATTGTTTTGGTG |
| SLIC T7 HnRNPDL-F | TGACAGGTGGCCAACAGATGGGTATGGAGGTCCCGCCCAGG |
| SLIC T7 HnRNPDL-R | GAGGGTCCATGGTGATACAAGGGATTAGTATGGCTGGTAATTGTTTTGGTG |
| TIAL1-F | ATGATGGAAGACGACGGGCAGC |
| TIAL1-R | TCACTGTGTTTGGTAACTTGCCATAC |
| SLIC T7 TIAL1-F | TGACAGGTGGCCAACAGATGGGTATGATGGAAGACGACGGGCAGC |
| SLIC T7 TIAL1-R | GAGGGTCCATGGTGATACAAGGGATCACTGTGTTTGGTAACTTGCCATAC |
| MAPT E3-12-F | ctggctaccaaagGTAATAAGAAGC |
| MAPT E3-12-R | ACCTTTGGTAGCCAGAAAAAAGGATGAGTG |
| MAPT E13-22-F | caaagGTGCAGATAATTTGGATCTTAGC |
| MAPT E13-22-R | AATTATCTGCACCTTTGGTAGCCAGAAAAAAGGATG |
| MAPT E23-32-F | GATAATTAATAAGAAGCCAACGTCCAG |
| MAPT E23-32-R | GCTTCTTATTAATTATCTGCACCTTTGGTAGCCAG |
| MAPT E33-42-F | GAAGCTGGATCTTAGTCCAAGTGTG |
| MAPT E33-42-R | CTAAGATCCAGCTTCTTATTAATTATCTGCACCTTTG |
| MAPT E43-52-F | CTTAGCAACGTCCAGGCTCAAAGGA |
| MAPT E43-52-R | CTGGACGTTGCTAAGATCCAGCTTCTTATT |
| MAPT E53-62-F | GTCCAGTCCAAGTGTGTAATATCAAACAC |
| MAPT E53-62-R | CACACTTGGACTGGACGTTGCTAAGATCCAGC |
| MAPT E63-72-F | GTGTGGCTCAAAGGACACGTCCCGGG |
| MAPT E63-72-R | TCCTTTGAGCCACACTTGGACTGGACGTTGC |
| MAPT E73-82-F | CAAAGGATAATATCAAAGAGGCGGCAGT |

| Name | Sequence (5′ to 3′) |
| --- | --- |
| MAPT E73-82-R | TTTGATATTATCCTTTGAGCCACACTTGGACTG |
| MAPT E83-91-F | TCAAACACGTCCCGGGTgtgagtaccttc |
| MAPT E83-91-R | CCGGGACGTGTTTGATATTATCCTTTGAGCC |
| MAPT IN9 1-10-F | ctcatcctttttctgGTGCAGATAAT |
| MAPT IN9 1-10-R | CAGAAAAAAGGATGAGTGACACGCCACCCTGGAC |
| MAPT IN9 11-20-F | ggcgtgtcactcatcgctaccaaagG |
| MAPT IN9 11-20-R | GATGAGTGACACGCCACCCTGGACCCGCCTGC |
| MAPT IN9 21-30-F | ggtccagggtggcgtctttttctg |
| MAPT IN9 21-30-R | ACGCCACCCTGGACCCGCCTGCTTGCTCGCAAG |
| MAPT IN9 31-40-F | caagcAggcgggtccgtcactcatc |
| MAPT IN9 31-40-R | GGACCCGCCTGCTTGCTCGCAAGGACGCCTCCAC |
| MAPT IN9 41-50-F | gtccttgcgagcaagcagggtggcgtg |
| MAPT IN9 41-50-R | GCTTGCTCGCAAGGACGCCTCCACTTTCGATGAGTGAC |
| MAPT IN10 1-10-F | CCGGGAGGCGGCAGTtcacacgtcc |
| MAPT IN10 1-10-R | ACTGCCGCCTCCCGGGACGTGTTTGATATTATC |
| MAPT IN10 11-20-F | GCAGTgtgagtacctcatgcgccgtgc |
| MAPT IN10 11-20-R | AGGTACTCACACTGCCGCCTCCCGGGACGTGTTTG |
| MAPT IN10 21-30-F | gtaccttcacacgtccgctgtggcttg |
| MAPT IN10 21-30-R | GGACGTGTGAAGGTACTCACACTGCCGCCTCC |
| MAPT IN10 31-40-F | cgtcccatgcgccgtgaattattag |
| MAPT IN10 31-40-R | ACGGCGCATGGGACGTGTGAAGGTACTCACAC |
| MAPT IN10 41-50-F | gccgtgctgtggcttgaagtggtgtg |
| MAPT IN10 41-50-R | AAGCCACAGCACGGCGCATGGGACGTGTGAAG |
| MAPT E10 63-68-F | GTGTGGCTCAAAGGACAAACACGTCCCG |
| MAPT E10 63-68-R | TCCTTTGAGCCACACTTGGACTGGACGTTGC |
| MAPT E10 7 0-74-F | CTCAAAGGATAATATCCGTCCCGGGAG |
| MAPT E170-74-R | GATATTATCCTTTGAGCCACACTTGGACTGGACGTTG |
| MAPT E10 62-68-F | AGTGTGGCTCAAAGGCAAACACGTCCCG |
| MAPT E10 62-68-R | CCTTTGAGCCACACTTGGACTGGACGTTG |
| MAPT E10 64-68-F | GTGTGGCTCAAAGGATCAAACACGTC |
| MAPT E10 64-68-R | ATCCTTTGAGCCACACTTGGACTGGACGTTG |
| MAPT E10 65-68-F | GTGGCTCAAAGGATACAAACACGTC |
| MAPT E10 65-68-R | TATCCTTTGAGCCACACTTGGACTGGACGTTG |
| MAPT E10 65-67-F | GTGGCTCAAAGGATATCAAACACGTC |
| MAPT E10 64-67-F | GTGTGGCTCAAAGGATTCAAACACGTC |
| MAPT E10 64-67-R | ATCCTTTGAGCCACACTTGGACTGGACGTTG |
| MAPT E10 64C-F | GTGTGGCTCAAAGGATCATATCAAACACGTC |
| MAPT E10 64C-R | ATCCTTTGAGCCACACTTGGACTGGACGTTG |
| MAPT E10 65C-F | GTGTGGCTCAAAGGATACTATCAAACACGTC |
| MAPT E10 65C-R | TATCCTTTGAGCCACACTTGGACTGGACGTTG |
| MAPT E10 64,65G-F | GTGTGGCTCAAAGGATCCTATCAAACACGTC |

| Name | Sequence (5′ to 3′) |
|---|---|
| MAPT E10 67C-F | GTGTGGCTCAAAGGATAATCTCAAACACGTC |
| MAPT E10 67C-R | ATTATCCTTTGAGCCACACTTGGACTGGACGTTG |
| MAPT E10 64,65,67C-F | GTGTGGCTCAAAGGATCCTCTCAAACACGTC |
| MAPT E10 66C-F | GTGTGGCTCAAAGGATAACATCAAACACGTC |
| MAPT E10 66C-R | TTATCCTTTGAGCCACACTTGGACTGGACGTTG |
| MAPT E10 68C-F | GTGGCTCAAAGGATAATACCAAACACGTC |
| MAPT E10 68C-R | TATTATCCTTTGAGCCACACTTGGACTGGACGTTG |
| MAPT E10 66,68C-F | GTGGCTCAAAGGATAACACCAAACACGTC |
| SLIC T7 λN AB-F | GCTCAATGGAAAGCTGCAAACATGTCGGAAGCGGGCGAG |
| SLIC T7 λN AB-R | CCAAACTCACCCTGAAGTTCTCATCAGTATGGCTTGTAGTTATTCTGATGG |
| T7λN-F | tgagaacttcagggtgagtttgg |
| T7λN-R | GTTTGCAGCTTTCCATTGAGC |
| Flag-F | TGAGTTTAAACCCGCTGATCAG |
| Flag-R | CTTGTCATCGTCATCCTTGTAG |
| SLIC Flag AB-F | CTACAAGGATGACGATGACAAGATGTCGGAAGCGGGCGAG |
| SLIC Flag AB-R | CTGATCAGCGGGTTTAAACTCATCAGTATGGCTTGTAGTTATTCTGATGG |
| RBMX/L1-F | ATGGTTGAAGCAGATCGCCCAG |
| RBMX/L1-R | CTAGTATCTGCTTCTGCCTCCC |
| SLIC-Flag-RBMX/L1-F | CTACAAGGATGACGATGACAAGATGGTTGAAGCAGATCGCCCAG |
| SLIC-Flag-RBMX/L1-R | CTGATCAGCGGGTTTAAACTCACTAGTATCTGCTTCTGCCTCCC |
| △CBFNT-F | AAAATGTTCGTTGGTGGC |
| △CBFNT-R | TCAGTATGGCTTGTAGTTATTCTGATGG |
| SLIC T7△CBFNT-F | TGACAGGTGGCCAACAGATGGGTAAAATGTTCGTTGGTGGC |
| SLIC T7△CBFNT-R | GAGGGTCCATGGTGATACAAGGGATCAGTATGGCTTGTAGTTATTCTGATGG |
| △RRMc-F | CAGCCCAAAGAAGTCTATC |
| △RRMc-R | GTAGCCCTGCTGGTAGCCG |
| SLIC T7△RRM-F | GAACGAGGAGGACGCGGGACAGCCCAAAGAAGTCTATC |
| SLIC T7△RRM-R | GTAGCCGCCATAGCCAGGCCCGTAGCCCTGCTGGTAGCCG |
| △RRM Vctr-F | GGGCCTGGCTATGGCGGCTAC |
| △RRM Vctr-R | TCCCGCGTCCTCCTCGTTC |
| △Gly-F | ATGTCGGAAGCGGGCGAG |

| Name | Sequence (5′ to 3′) |
|---|---|
| △Gly-R | GGCCACCTTGATCTCACAC |
| SLIC T7△Gly-F | TGACAGGTGGCCAACAGATGGGTATGTCGGAAGCGGGCGAG |
| SLIC T7△Gly-R | GAGGGTCCATGGTGATACAAGGGAGGCCACCTTGATCTCACAC |
| SLIC FlAG△CBFNT-F | CTACAAGGATGACGATGACAAGAAAATGTTCGTTGGTGGC |
| SLIC FlAG△CBFNT-R | CTGATCAGCGGGTTTAAACTCATCAGTATGGCTTGTAGTTATTCTGATGG |
| △RRM-F | ATGTCGGAAGCGGGCGAG |
| △RRM-R | TCAGTATGGCTTGTAGTTATTCTGATGG |
| SLIC FLAG△RRM-F | CTACAAGGATGACGATGACAAGATGTCGGAAGCGGGCGAG |
| SLIC FLAG△Gly-R | CTGATCAGCGGGTTTAAACTCAGGCCACCTTGATCTCACAC |

**Table S2**: **Sequences of ASOs used in this study.**

| Name | Sequence (5′ to 3′) |
|---|---|
| ASO-1 | TCCAACTCTGACTCTGACct |
| ASO-2 | CCTGATTCCAACTCTGACTC |
| ASO-3 | CGTAGCCCTGATTCCAACTC |
| ASO-4 | AGTTGCCGTAGCCCTGATTC |
| ASO-5 | TCCAGTAGTTGCCGTAGCCC |
| ASO-6 | CCTGGTTCCAGTAGTTGCCG |
| ASO-7 | CGTAGCCCTGGTTCCAGTAG |
| ASO-8 | GGTAGCCGTAGCCCTGGTTC |
| ASO-9 | CCTGCTGGTAGCCGTAGCCC |
| ASO-10 | CGTAGCCCTGCTGGTAGCCG |
| ASO-11 | CAGGCCCGTAGCCCTGCTGG |
| ASO-12 | CATAGCCAGGCCCGTAGCCC |
| ASO-13 | AGCCGCCATAGCCAGGCCCG |
| ASO-14 | AGTCGTAGCCGCCATAGCCA |
| ASO-15 | GCGAGTAGTCGTAGCCGCCA |
| ASO-16 | CATAGGGCGAGTAGTCGTAG |
| ASO-17 | AATAGCCATAGGGCGAGTAG |
| ASO-18 | AGCCGTAATAGCCATAGGGC |
| ASO-19 | GGCCGTAGCCGTAATAGCCA |
| ASO-20 | AGCCGGGGCCGTAGCCGTAA |
| ASO-21 | AGTCGTAGCCGGGGCCGTAG |
| ASO-22 | tacTGTAGTCGTAGCCGGGG |
| ASO-23 | TCTGACTCTGACctgtgggg |
| ASO-24 | TCTGACctgtgggggagca |
| ASO-25 | ctgtgggggagcagggcac |
| ASO-26 | ggggagcagggcacaggggc |
| ASO-27 | cagggcacaggggcccgtgg |
| ASO-28 | acaggggcccgtggaccatt |
| ASO-29 | gcccgtggaccatttagcat |
| ASO-30 | ggaccatttagcatacgcag |
| ASO-31 | tttagcatacgcaggagcta |
| ASO-32 | atacgcaggagctaggatgg |
| ASO-33 | cctacttacTGTAGTCGTAG |
| ASO-34 | ctctctcctacttacTGTAG |
| ASO-35 | gcctccctctctcctactta |
| ASO-36 | gatggggcctccctctctcc |
| ASO-37 | tgagcggatggggcctccct |
| ASO-38 | gaggggtgagcggatggggc |
| ASO-39 | ggggacgaggggtgagcgga |
| ASO-40 | tcccctggggacgaggggtg |
| ASO-41 | cctgcctcccctggggacga |

| Name | Sequence (5′ to 3′) |
|---|---|
| ASO-42 | cactgtcctgcctcccctgg |
| ASO-00 | CACCTTTGATACAACTACCG |
| ASO-NC | GAGTTACCCGACACGTATGA |

**Table S3: Primer sequences used for qRT-PCR and RT-PCR.**

| Name | Sequence (5′ to 3′) |
|---|---|
| hnRNPAB-F | TGTGAGATCAAGGTGGCCCAG |
| hnRNPAB-R | TATCCAAACAAAGCATGTGTGCG |
| APOER2-F | TGGTGATAGCCCTCCTGTG |
| APOER2-R | TGCATGGGACTGAATTCC |
| TMPO-F | ACTCTAAAATAGAGCTCAAGCTTG |
| TMPO-R | TTCAGCTCTTGTCAATGGTTTC |
| ZDHHC7-F | TGGCTGACCGGGTCTGGTTC |
| ZDHHC7-R | GGGCGCGCTCGGGTTTAATAC |
| MCL-1-F | GAGGAGGAGGAGGACGAGTT |
| MCL-1-R | AACCAGCTCCTACTCCAGCA |
| STAG2-F | CTGAAGAAAGTAGTAGTAGTGACAG |
| STAG2-R | CCATGGTGTCAAAATCCATTC |
| PICALM-F | CACATCAAAGCTGCCCAATGATC |
| PICALM-R | CCCCAGAATCTACAATAACATTTG |
| NT5E-F | GCTCAGAAAGTGAGGGGTGT |
| NT5E-R | TGGAAGGTGGATTGCCTGTG |
| NR1D1-F | CTTGTCTCTGCAGACCGCT |
| NR1D1-R | GCTTTTCCTTTTCGTCTCGTAA |
| GAPDH-F | TCAACGACCACTTTGTCAAGCTCA |
| GAPDH-R | GCTGGTGGTCCAGGGGTCTTACT |

**Table S4: Antibodies used in this study.**

| Name | SOURCE | IDENTIFIER |
|---|---|---|
| Rabbit polyclonal anti-hnRNPAB | ABclonal | A17497 |
| Rabbit monoclonal anti- hnRNPAB | Abcam | ab199724 |
| Rabbit monoclonal anti-T7 | Abcam | ab317258 |
| Rabbit polyclonal anti-hnRNPDL | Abcam | ab272661 |
| Rabbit monoclonal anti-TIAL1 | Abcam | ab169547 |
| Rabbit monoclonal anti-RBMX | Abcam | Ab190352 |
| Rabbit polyclonal anti-RBMXL1 | Invitrogen | PA5-71058 |
| Mouse monoclonal anti-FLAG | Proteintech | 66008-4-Ig |
| Rabbit polyclonal anti-RPL4 | ABclonal | A5886 |
| Rabbit polyclonal anti-TUFM | ABclonal | A6423 |
| Rabbit Monoclonal anti-HADHB | ABclonal | A25207 |
| Rabbit polyclonal anti-EIF2S3 | ABclonal | A6581 |
| Rabbit Monoclonal anti-EIF4A1 | Abcam | ab185946 |
| Rabbit Monoclonal anti-4R-tau | Cell Signaling Technology | 79327S |
| Rabbit Monoclonal anti-4R-tau | Abcam | ab218314 |
| Mouse monoclonal anti-AT8 | Invitrogen | MN1020 |
| IRDye 680RD Goat anti-Rabbit IgG Secondary Antibody | LI-COR Biosciences | 926-68071 |
| IRDye 680RD Goat anti-Mouse IgG Secondary Antibody | LI-COR Biosciences | 926-68070 |
